# C-type natriuretic peptide facilitates autonomic Ca^2+^ entry in growth plate chondrocytes for stimulating bone growth

**DOI:** 10.1101/2021.09.17.460760

**Authors:** Yuu Miyazaki, Atsuhiko Ichimura, Ryo Kitayama, Naoki Okamoto, Tomoki Yasue, Feng Liu, Yohei Ueda, Ichiro Yamauchi, Takuro Hakata, Kazumasa Nakao, Sho Kakizawa, Miyuki Nishi, Yasuo Mori, Haruhiko Akiyama, Kazuwa Nakao, Hiroshi Takeshima

## Abstract

The growth plates are cartilage tissues found at both ends of developing bones, and vital proliferation and differentiation of growth plate chondrocytes are primarily responsible for bone growth. C-type natriuretic peptide (CNP) stimulates bone growth by activating natriuretic peptide receptor 2 (NPR2) which is equipped with guanylate cyclase on the cytoplasmic side, but its signaling pathway is unclear in growth plate chondrocytes. We previously reported that transient receptor potential melastatin-like 7 (TRPM7) channels mediate intermissive Ca^2+^ influx in growth plate chondrocytes, leading to activation of Ca^2+^/calmodulin-dependent protein kinase II (CaMKII) for promoting bone growth. In this report, we provide experimental evidence indicating a functional link between CNP and TRPM7 channels. Our pharmacological data suggest that CNP-evoked NPR2 activation elevates cellular cGMP content and stimulates big-conductance Ca^2+^-dependent K^+^ (BK) channels as a substrate for cGMP-dependent protein kinase (PKG). BK channel-induced hyperpolarization likely enhances the driving force of TRPM7-mediated Ca^2+^ entry and seems to accordingly activate CaMKII. Indeed, *ex vivo* organ culture analysis indicates that CNP-facilitated bone growth is abolished by chondrocyte-specific *Trpm7* gene ablation. The defined CNP signaling pathway, the NPR2-PKG-BK channel-TRPM7 channel-CaMKII axis, likely pinpoints promising target proteins for developing new therapeutic treatments for divergent growth disorders.

## Introduction

The development of skeletal long bones occurs through endochondral ossification processes, during which chondrocyte layers form the growth plates at both ends of bone rudiments, and then the expanded cartilage portions are gradually replaced by trabecular bones through the action of osteoclasts and osteoblasts [1]. Therefore, bone size largely depends on the proliferation of growth plate chondrocytes during endochondral development. On the other hand, atrial (ANP), brain (BNP) and C-type (CNP) natriuretic peptides regulate diverse cellular functions by activating the receptor guanylate cyclases, NPR1 and NPR2 [2]. Of the natriuretic peptides, CNP exclusively stimulates bone development by acting on growth plate chondrocytes expressing the CNP-specific receptor NPR2 [2–4]. Indeed, loss- and gain-of-function mutations in the human *NPR2* gene cause acromesomelic dysplasia and skeletal overgrowth disorder, respectively [5, 6]. Furthermore, translational studies have been probing the benefits of CNP treatments in various animal models with impaired skeletal growth, and a phase III clinical trial of CNP therapy has recently started in patients with chondrodystrophia primarily resulting from mutations in the *FGFR3* gene [7]. It is thus likely that NPR2 guanylate cyclase controls chondrocytic cGMP content during growth plate development. Downstream of NPR2 activation, cGMP-dependent protein kinase (PKG) seems to phosphorylate target proteins to facilitate growth plate chondrogenesis [4]. Activated PKG is postulated to stimulate the biosynthesis of growth plate extracellular matrix by playing an inhibitory role in the mitogen-activated protein kinase Raf-MEK-ERK cascade [8]. In parallel, glycogen synthase kinase 3β (GSK3β) is likely activated by PKG-mediated phosphorylation, leading to the hypertrophic maturation of growth plate chondrocytes [9]. However, it is still unclear how CNP promotes bone growth at the molecular level, and it is important to further address CNP signaling cascade in growth plate chondrocytes.

In the transient receptor potential channel superfamily, the melastatin subfamily member 7 (TRPM7) forms a mono- and divalent cation-permeable channel in various cell types and participates in important cellular processes including cell growth and adhesion [10]. We recently reported that growth plate chondrocytes generate autonomic intracellular Ca^2+^ fluctuations, which are generated by the intermittent gating of TRPM7 channels, and also that TRPM7-mediated Ca^2+^ entry activates Ca^2+^/calmodulin-dependent protein kinase II (CaMKII), facilitating chondrogenesis for endochondral bone development [11]. Based on these observations, we explored the link between CNP signaling and TRPM7-mediated Ca^2+^ entry through the experiments described in this report. Our data obtained clearly indicate that big-conductance Ca^2+^-dependent K^+^ (BK) channels play a key role in the functional coupling between NPR2 and TRPM7 channels in growth plate chondrocytes.

## Results

### CNP facilitates spontaneous Ca^2+^ fluctuations in growth plate chondrocytes

In the growth plates of developing bones, proliferating cartilage cells, designated as round and columnar chondrocytes, frequently exhibit weak increases and decreases in intracellular Ca^2+^ concentration under resting conditions [11]. On the other hand, previous *in vivo* studies demonstrated that CNP application (>1 μmol/kg) stimulates endochondral bone growth [2]. In our Fura-2 imaging of round chondrocytes within femoral bone slices prepared from wild-type mice, CNP pretreatments (30∼300 nM for 1 hr) dose-dependently facilitated spontaneous Ca^2+^ fluctuations (Figure 1A). In particular, fluctuation-positive cell ratio and fluctuation amplitude were remarkably elevated in response to the CNP treatments. In contrast, ANP treatments exerted no effects on Ca^2+^ fluctuations in growth plate chondrocytes.

**Figure 1.**
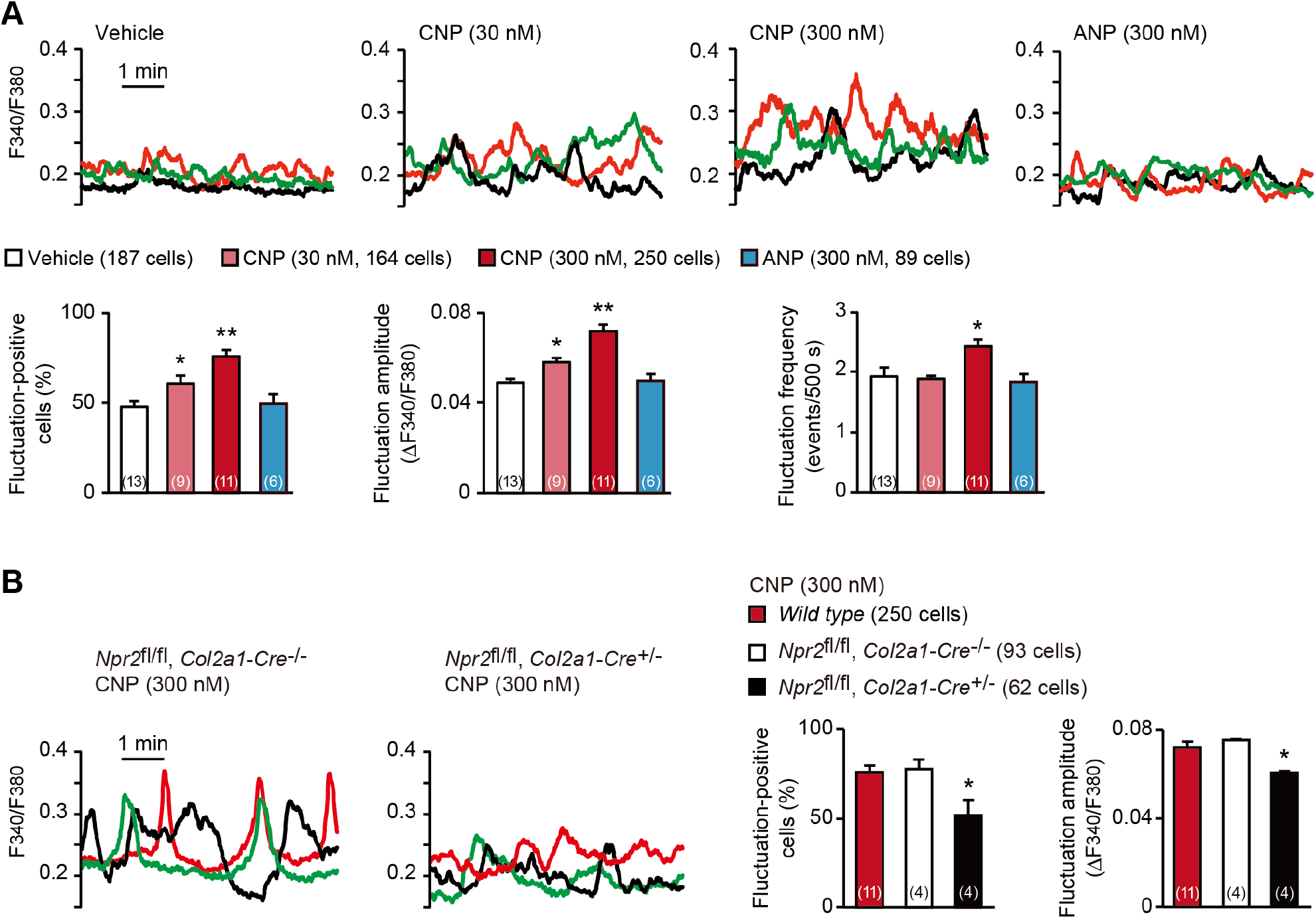
CNP-induced facilitation of Ca^2+^ fluctuations in growth plate chondrocytes. **(A)** Fura-2 imaging of round chondrocytes pretreated with or without natriuretic peptides. Femoral bone slices prepared from wild-type C57BL embryos were pretreated with or without CNP and ANP, and subjected to Ca^2+^ imaging. Representative recording traces from three cells are shown in each pretreatment group (upper panels). The effects of CNP and ANP pretreatments on spontaneous Ca^2+^ fluctuations are summarized (lower graphs). The fluctuation-positive cell ratio, fluctuation amplitude and frequency were statistically analyzed, and significant differences from the control vehicle pretreatment are marked with asterisks (\**p*<0.05 and \*\**p*<0.01 in one-way ANOVA and Dunnett’s test). The data are presented as the means ± SEM. with *n* values indicating the number of examined mice. **(B)** Fura-2 imaging of round chondrocytes prepared from chondrocyte-specific *Npr2*-knockout (*Npr2*^fl/fl^, *Col2a1-Cre*^+/-^) and control (*Npr2*^fl/fl^, *Col2a1-Cre*^-/-^) mice. The bone slices were pretreated with CNP, and then subjected to Ca^2+^ imaging. Representative recording traces are shown (left panels) and the CNP-pretreated effects are summarized (right graphs); significant differences from the wild-type group are marked with asterisks (\**p*<0.05 in one-way ANOVA and Tukey’s test). The data are presented as the means ± SEM. with *n* values indicating the number of examined mice.

In chondrocyte-specific *Npr2*-knockout mice (*Npr2*^fl/fl^, *Col2a1-Cre*^+/−^), Cre recombinase is expressed under the control of the collagen type 2α1 gene promoter and thus inactivates the floxed *Npr2* alleles in a chondrocyte-specific manner [12]. Our RT-PCR analysis indicated that the floxed *Npr2* gene was largely inactivated in the growth plates prepared from the E17.5 mutant embryos, but such recombination events were not detected in other tissues examined (Figure 1-figure supplement 1A and B). Accordingly, *Npr2* mRNA contents in the mutant growth plates were reduced to less than 40% of controls (Figure 1-figure supplement 1C), despite the growth plate preparations contain not only chondrocytes but also perichondrium-resident cells including undifferentiated mesenchymal cells and immature chondroblasts. In contrast to the imaging observations in wild-type and control bone slices, CNP treatments failed to enhance Ca^2+^ fluctuations in the mutant round chondrocytes prepared from the chondrocyte-specific *Npr2*-knockout mice (Figure 1B). Therefore, CNP seems to facilitate spontaneous Ca^2+^ fluctuations downstream of NPR2 activation in growth plate chondrocytes.

### Activated PKG facilitates spontaneous Ca^2+^ fluctuations

CNP binds to NPR2 to activate its intrinsic guanylate cyclase and thus stimulates PKG by elevating cellular cGMP contents [2]. CNP also binds to NPR3 which acts as a decoy receptor for ligand clearance, but the *Npr3* gene seemed to be inactive in growth plate chondrocytes (Figure 1-figure supplement 2). Next, we pharmacologically verified the contribution of PKG to CNP-facilitated Ca^2+^ fluctuations. The cGMP analog 8-(4-chlorophenylthio)-cyclic GMP (8-pCPT-cGMP) is widely used as a PKG-selective activator, while KT5823 is a typical PKG inhibitor. In wild-type growth plate chondrocytes pretreated with 8-pCPT-cGMP (100 μM for 1 hr), spontaneous Ca^2+^ fluctuations were remarkably facilitated (Figure 2A); both fluctuation-positive cell rate and fluctuation amplitude were highly increased. In contrast, the bath application of KT5823 (2 μM) clearly attenuated CNP-facilitated Ca^2+^ fluctuations within a short time frame (Figure 2B). Therefore, PKG activation seems to be essential for CNP-facilitated Ca^2+^ fluctuations in growth plate chondrocytes.

**Figure 2.**
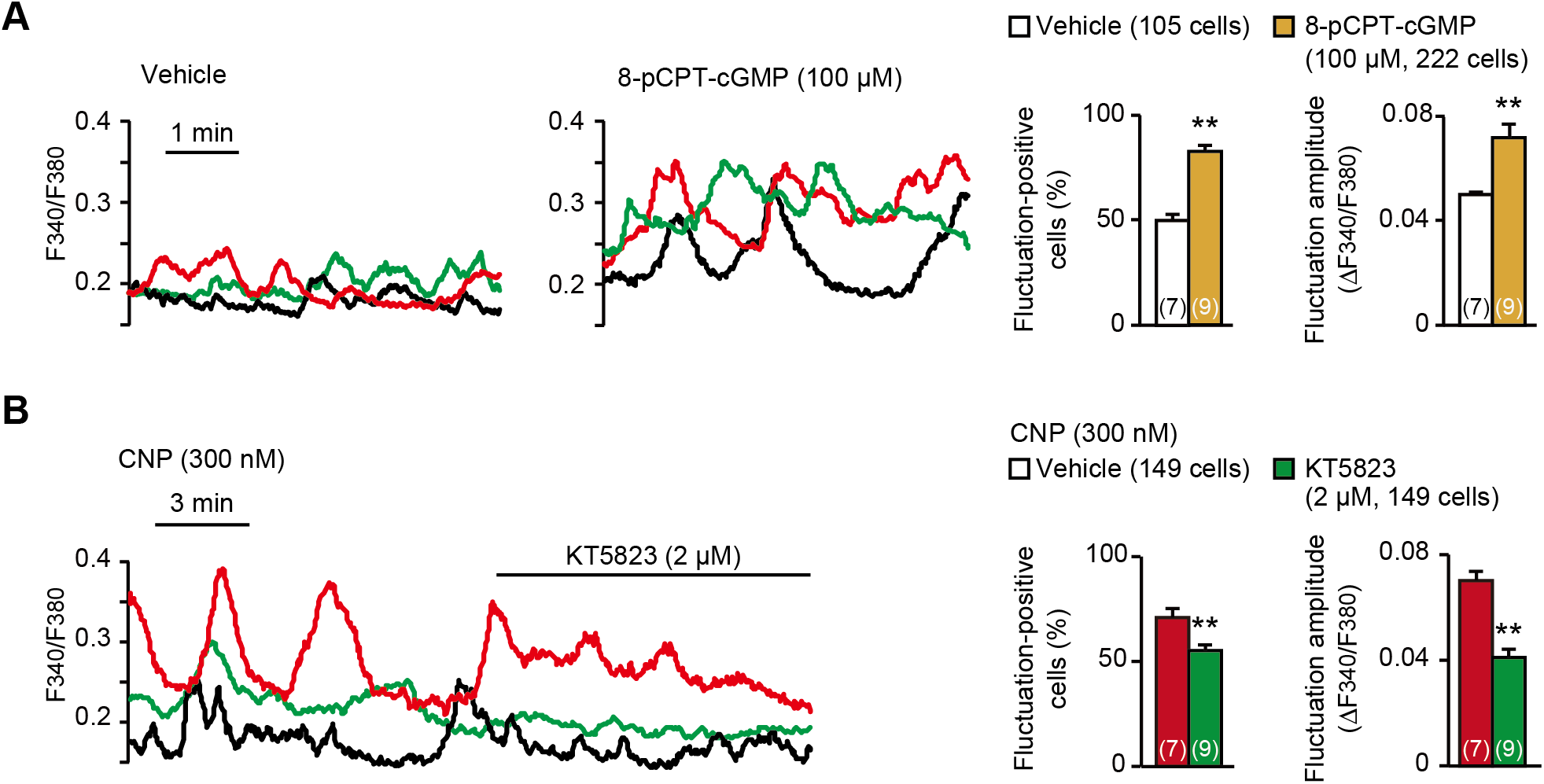
Contribution of PKG to CNP-facilitated Ca^2+^ fluctuations. **(A)** Facilitated Ca^2+^ fluctuations in round chondrocytes pretreated with the PKG activator 8-pCPT-cGMP. Wild-type bone slices were pretreated with or without the cGMP analog, and then subjected to Ca^2+^ imaging. Representative recording traces are shown (left panels), and the pharmacological effects are summarized (right graphs). Significant differences between control and 8-pCPT-cGMP pretreatments are marked with asterisks (\*\**p*<0.01 in *t*-test). The data are presented as the means ± SEM. with *n* values indicating the number of examined mice. **(B)** Attenuation of CNP-facilitated Ca^2+^ fluctuations by the PKG inhibitor KT5823. Wild-type bone slices were pretreated with CNP, and then subjected to Ca^2+^ imaging. Representative recording traces are shown (left panel), and KT5823-induced effects are summarized (right graphs). Significant KT5823-induced shifts are marked with asterisks (\*\**p*<0.01 in *t*-test). The data are presented as the means ± SEM. with *n* values indicating the number of examined mice.

### Activated BK channels contribute to CNP-facilitated Ca^2+^ fluctuations

Spontaneous Ca^2+^ fluctuations are facilitated by activated BK channels in growth plate chondrocytes [11]. Previous studies have established a functional link between PKG and BK channels in several cell types including smooth muscle and endothelial cells; activated PKG enhances BK channel gating by directly phosphorylating the α subunit KCNMA1 protein [13–15]. We thus examined whether altered BK channel activity is associated with CNP-facilitated Ca^2+^ fluctuations. The BK channel inhibitor paxilline (10 μM) exerted no obvious effects on basal Ca^2+^ fluctuations in non-treated chondrocytes. However, the same paxilline treatments remarkably inhibited CNP-facilitated Ca^2+^ fluctuations (Figure 3A); both fluctuation-positive cell ratio and fluctuation amplitude were clearly decreased after paxilline application. On the other hand, the BK channel activator NS1619 (30 μM) stimulated basal Ca^2+^ fluctuations in the growth plate chondrocytes prepared from control mice. The NS1619-induced effects were preserved in the mutant chondrocytes prepared from chondrocyte-specific *Npr2*-knockout mice (Figure 3B). Therefore, BK channel activation is likely involved in CNP-facilitated Ca^2+^ fluctuations in growth plate chondrocytes.

**Figure 3.**
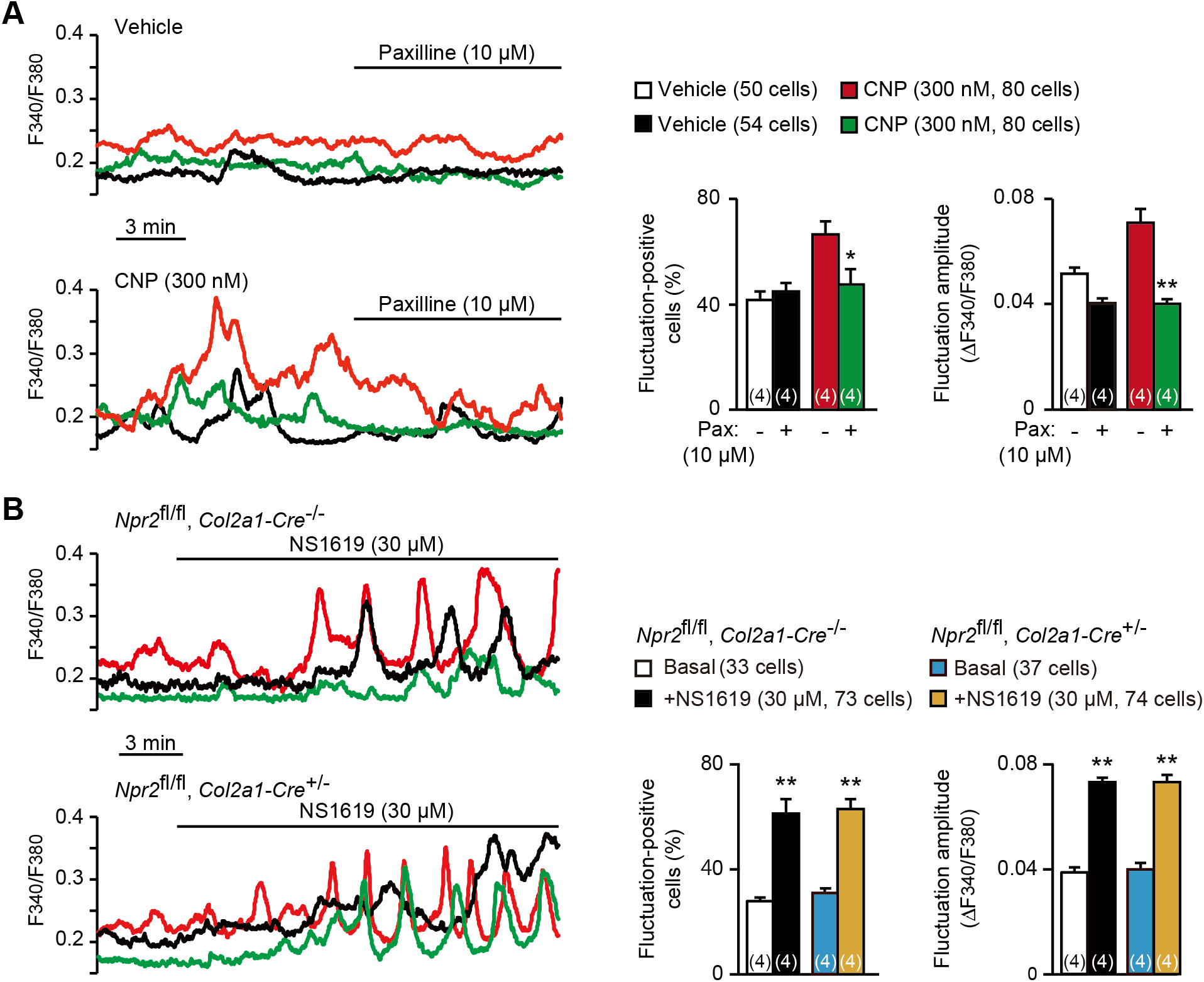
Contribution of BK channels to CNP-facilitated Ca^2+^ fluctuations. **(A)** Attenuation of CNP-facilitated Ca^2+^ fluctuations by the BK channel inhibitor paxilline in round chondrocytes. Wild-type bone slices were pretreated with or without CNP, and then subjected to Ca^2+^ imaging. Representative recording traces are shown (left panels), and paxilline-induced effects are summarized (right graphs). Significant paxilline-induced shifts are marked with asterisks (\**p*<0.05 and \*\**p*<0.01 in one-way ANOVA and Tukey’s test). The data are presented as the means ± SEM. with *n* values indicating the number of examined mice. **(B)** Ca^2+^ fluctuations facilitated by the BK channel activator NS1619 in *Npr2*-deficient chondrocytes. Bone slices were prepared from the chondrocyte-specific *Npr2*-knockout and control embryos, and NS1619-induced effects were examined in Ca^2+^ imaging. Representative recording traces are shown (left panels), and the effects of NS1619 are summarized (right graphs). Significant NS1619-induced shifts are marked with asterisks (\*\**p* <0.01 in one-way ANOVA and Tukey’s test). The data are presented as the means ± SEM. with *n* values indicating the number of examined mice.

### PLC seems unrelated to CNP-facilitated Ca^2+^ fluctuations

Ca^2+^ fluctuations are maintained by phosphatidylinositol (PI) turnover in growth plate chondrocytes [11]. Although it has been reported that activated PKG inhibits phospholipase C (PLC) in smooth muscle [16–19], it might be possible that NPR2 activation enhances basal PLC activity to facilitate Ca^2+^ fluctuations. The PLC inhibitor U73122 (10 μM) remarkably inhibited basal Ca^2+^ fluctuations in non-treated chondrocytes: the fluctuation-positive cell ratio and fluctuation amplitude reduced less than half in response to U73122 application (Figure 3-figure supplement 3). U73122 was also effective for CNP-facilitated Ca^2+^ fluctuations, but the inhibitory efficiency seemed relatively weak compared to those on basal fluctuations. Given the different inhibitory effects, it is rather unlikely that PLC activation accompanies CNP-facilitated Ca^2+^ fluctuations.

PKG stimulates sarco/endoplasmic reticulum Ca^2+^-ATPase (SERCA) by phosphorylating the Ca^2+^ pump regulatory peptide phospholamban (PLN) in smooth and cardiac muscle cells [20–22], and activated Ca^2+^ pumps generally elevate stored Ca^2+^ content and thus stimulate store Ca^2+^ release. RT-PCR data suggested that the *Pln* gene and the *Atp2a2* gene encoding SERCA2 are weakly active in growth plate chondrocytes (Figure 1-figure supplement 2). To examine the effects of CNP treatments on Ca^2+^ stores, we examined Ca^2+^ responses to the activation of Gq-coupled lysophosphatidic acid (LPA) receptors (Figure 3-figure supplement 4A) and the Ca^2+^ pump inhibitor thapsigargin (Figure 3-figure supplement 4B). CNP- and vehicle-pretreated chondrocytes exhibited similar LPA-induced Ca^2+^ release and thapsigargin-induced Ca^2+^ leak responses. Therefore, CNP treatments seem ineffective for store Ca^2+^ pumps in growth plate chondrocytes. Moreover, the dose-dependency of Ca^2+^ release by LPA (1∼10 μM) was not altered between CNP- and vehicle-pretreated chondrocytes, implying that CNP does not affect basal PLC activity.

Among diverse Ca^2+^ handling-related proteins, PLC, PLN and BK channels have been reported as PKG substrates, however, our observations suggested that both PLC and PLN receive no obvious functional regulation in CNP-treated chondrocytes. On the other hand, the paxilline treatments diminished CNP-facilitated Ca^2+^ fluctuations down to non-treated control levels (Figure 3A), suggesting that activated BK channels predominantly contribute to CNP-facilitated Ca^2+^ fluctuations in growth plate chondrocytes.

### CNP induces BK channel-mediated hyperpolarization

To confirm the contribution of activated BK channels to CNP-facilitated Ca^2+^ fluctuations, we conducted confocal imaging using the voltage-dependent dye oxonol VI. In this imaging analysis, depolarization results in the accumulation of the dye into cells, in which the fractional fluorescence intensity, normalized to the maximum intensity monitored in the bath solution containing 100 mM KCl, is thus increased (Figure 4A left panel). The fractional intensity of CNP-pretreated cells was significantly lower than that of non-treated cells in a normal bath solution (Figure 4A middle graph), although both cells exhibited similar intensity shifts in high K^+^ bath solutions. Based on the recording data, we prepared a calibration plot for the relationship between the fractional intensity and theoretical membrane potential (Figure 4A right panel). In the tentative linear correlation, resting potentials of -46.4 ± 0.2 and -43.6 ± 0.3 mV were estimated in CNP-treated and non-treated cells, respectively. The estimated potentials closely approximate the reported value from monitoring articular chondrocytes using sharp microelectrodes [23].

**Figure 4.**
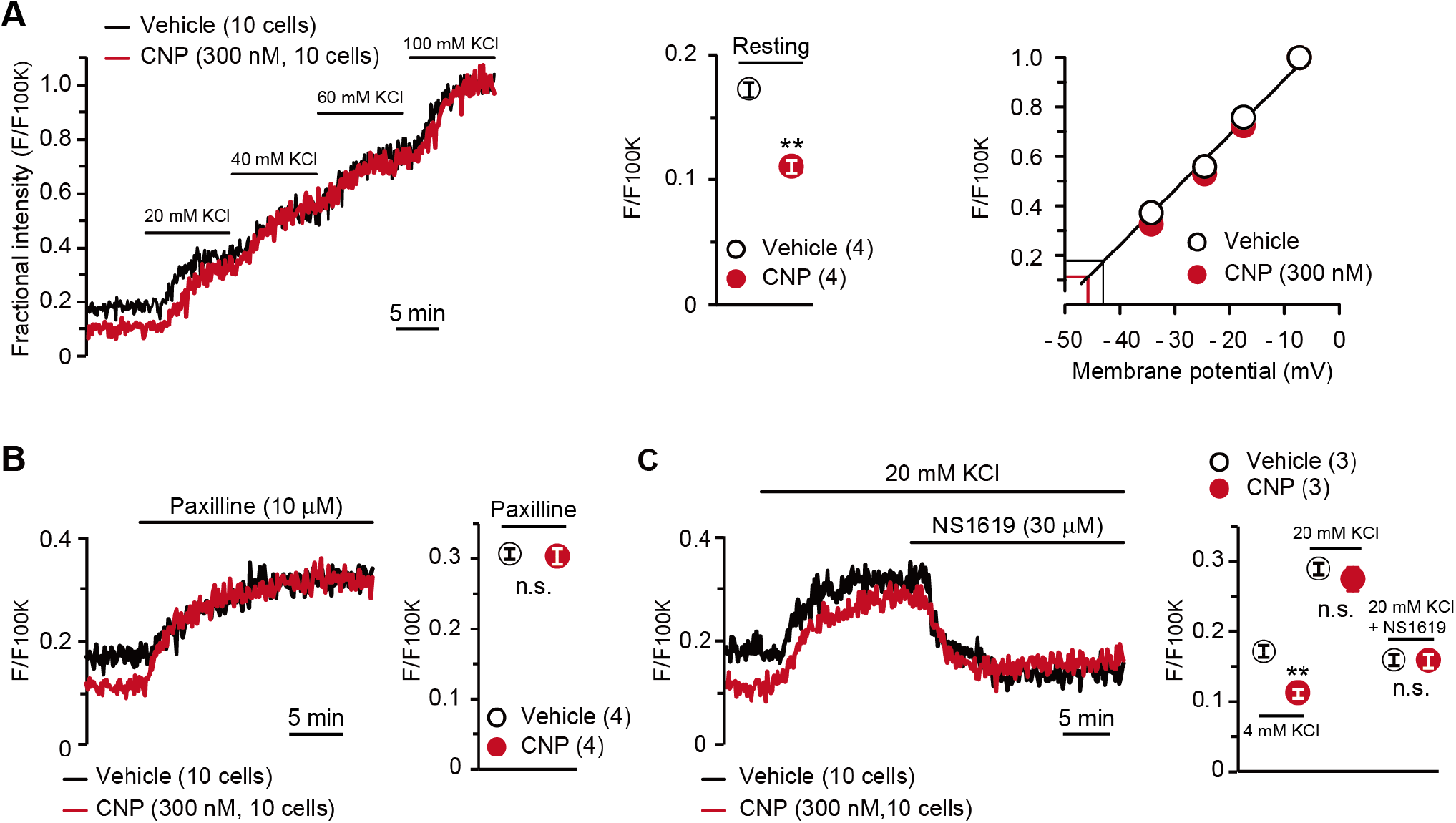
BK channel-mediated hyperpolarization induced by CNP. **(A)** Oxonol VI imaging of round chondrocytes pretreated with or without CNP. Wild-type bone slices were pretreated with or without CNP, and then subjected to membrane potential imaging. During contiguous treatments with high-K^+^ solutions, cellular fluorescence intensities were monitored and normalized to the maximum value in the 100 mM KCl-containing solution to yield the fractional intensity (left panel). The resting fractional intensities were quantified and statistically analyzed in CNP- and vehicle-pretreated cells (middle graph). For preparing the calibration plot (right panel), the data from ten cells in bathing solutions containing 4 (normal solution), 20, 40, 60 and 100 mM KCl are summarized; red and black lines indicate the estimated resting membrane potentials of CNP- and vehicle-pretreated cells, respectively. **(B)** Effects of the BK channel inhibitor paxilline on resting membrane potential in round chondrocytes. Recording data from ten cells pretreated with or without CNP were averaged (left panel), and the fractional intensities elevated by paxilline are summarized (right graph). **(C)** Effects of the BK channel activator NS1619 on membrane potential in round chondrocytes. Recording data from ten cells pretreated with or without CNP were averaged (left panel), and the fractional intensities in normal, 20 mM KCl and NS1619-containing 20 mM KCl solutions are summarized (right graph). Significant differences between CNP- and vehicle-pretreated cells are indicated by asterisks in **A** (\*\**p*<0.01 in *t*-test) and in **C** (\*\**p*<0.01 in one-way ANOVA and Dunn’s test). The data are presented as the means ± SEM. with *n* values indicating the number of examined mice.

In pharmacological assessments, paxilline elevated fractional intensities to the same levels in CNP-and non-treated chondrocytes (Figure 4B). Moreover, NS1619 decreased fractional intensities to the same levels in both cells under 20 mM KCl bathing conditions, which enabled us to reliably evaluate the reducing intensity shifts (Figure 4C). The oxonol VI imaging data suggested that CNP treatments induce BK channel-mediated hyperpolarization and thus facilitate spontaneous Ca^2+^ fluctuations by enhancing Ca^2+^-driving forces in growth plate chondrocytes.

### CNP enhances TRPM7-mediated Ca^2+^ entry and CaMKII activity

Spontaneous Ca^2+^ fluctuations are predominantly attributed to the intermissive gating of cell-surface TRPM7 channels in growth plate chondrocytes [11]. For pharmacological characterization of TRPM7 channels, FTY720 is used as a typical inhibitor, while NNC550396 is an activator. As reasonably expected, bath application of FTY720 (10 μM) clearly diminished CNP-facilitated Ca^2+^ fluctuations in round chondrocytes (Figure 5A). On the other hand, NNC550396 (30 μM) remarkably facilitated Ca^2+^ fluctuation in non-treated chondrocytes, and this facilitation was preserved in the mutant chondrocytes prepared from chondrocyte-specific *Npr2*-knockout mice (Figure 5B). Therefore, CNP treatments likely facilitate TRPM7-mediated Ca^2+^ influx in growth plate chondrocytes.

**Figure 5.**
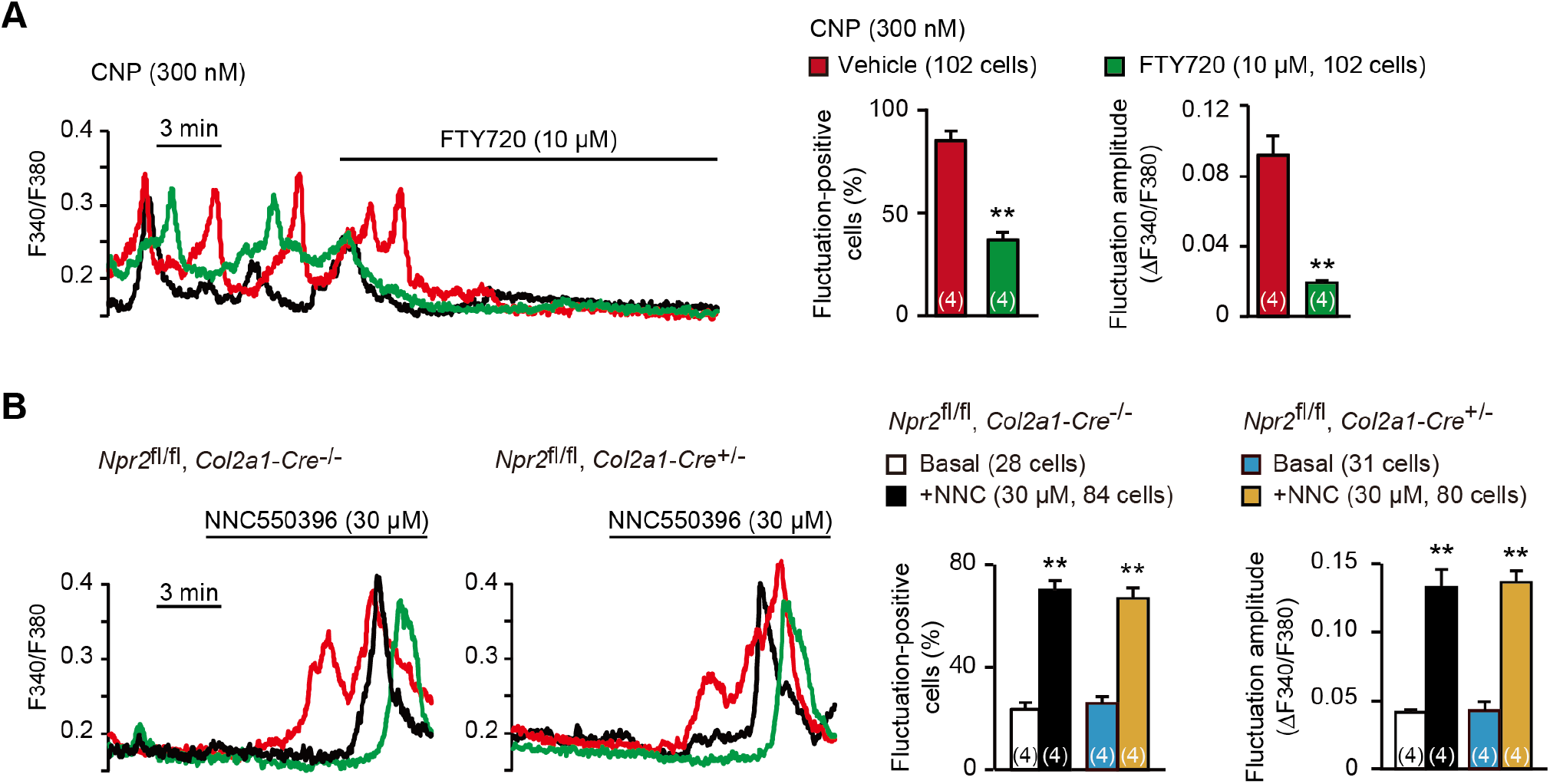
Enhanced TRPM7-mediated Ca^2+^ entry by CNP treatments. **(A)** Inhibition of CNP-facilitated Ca^2+^ fluctuations by the TRPM7 inhibitor FTY720 in round chondrocytes. Wild-type bone slices were pretreated with CNP, and then subjected to Ca^2+^ imaging. Representative recording traces are shown (left panel), and the effects of FTY720 are summarized (right graphs). Significant FTY720-induced shifts are marked with asterisks (\*\**p*<0.01 in *t*-test). The data are presented as the means ± SEM. with *n* values indicating the number of examined mice. **(B)** Ca^2+^ fluctuations facilitated by the TRPM7 channel activator NNC550396 in *Npr2*-deficient round chondrocytes. Bone slices were prepared from the chondrocyte-specific *Npr2*-knockout and control embryos, and NNC550396-induced effects were examined in Ca^2+^ imaging. Representative recording traces are shown (left panels) and the effects of NNC550396 on Ca^2+^ fluctuations are summarized (right graphs). Significant NNC550396-induced shifts in each genotype are marked with asterisks (\*\**p* <0.01 in one-way ANOVA and Tukey’s test). The data are presented as the means ± SEM. with *n* values indicating the number of examined mice.

TRPM7-mediated Ca^2+^ entry activates CaMKII in growth plate chondrocytes toward bone outgrowth [11], and cellular CaMKII activity can be estimated by immunochemically quantifying its autophosphorylated form. In immunocytochemical analysis, CNP-pretreated growth plate chondrocytes were more decorated with the antibody against phospho-CaMKII than non-treated control cells (Figure 6A). This CNP-facilitated decoration was abolished by the cotreatment of the CaMKII inhibitor KN93 (30 μM). This observation was further confirmed by Western blot analysis; CNP treatments increased the phospho-CaMKII population without affecting total CaMKII content in the cell lysates prepared from growth plates (Figure 6B). Therefore, CaMKII is likely activated downstream of enhanced TRPM7-mediated Ca^2+^ entry in CNP-treated growth plate chondrocytes.

**Figure 6.**
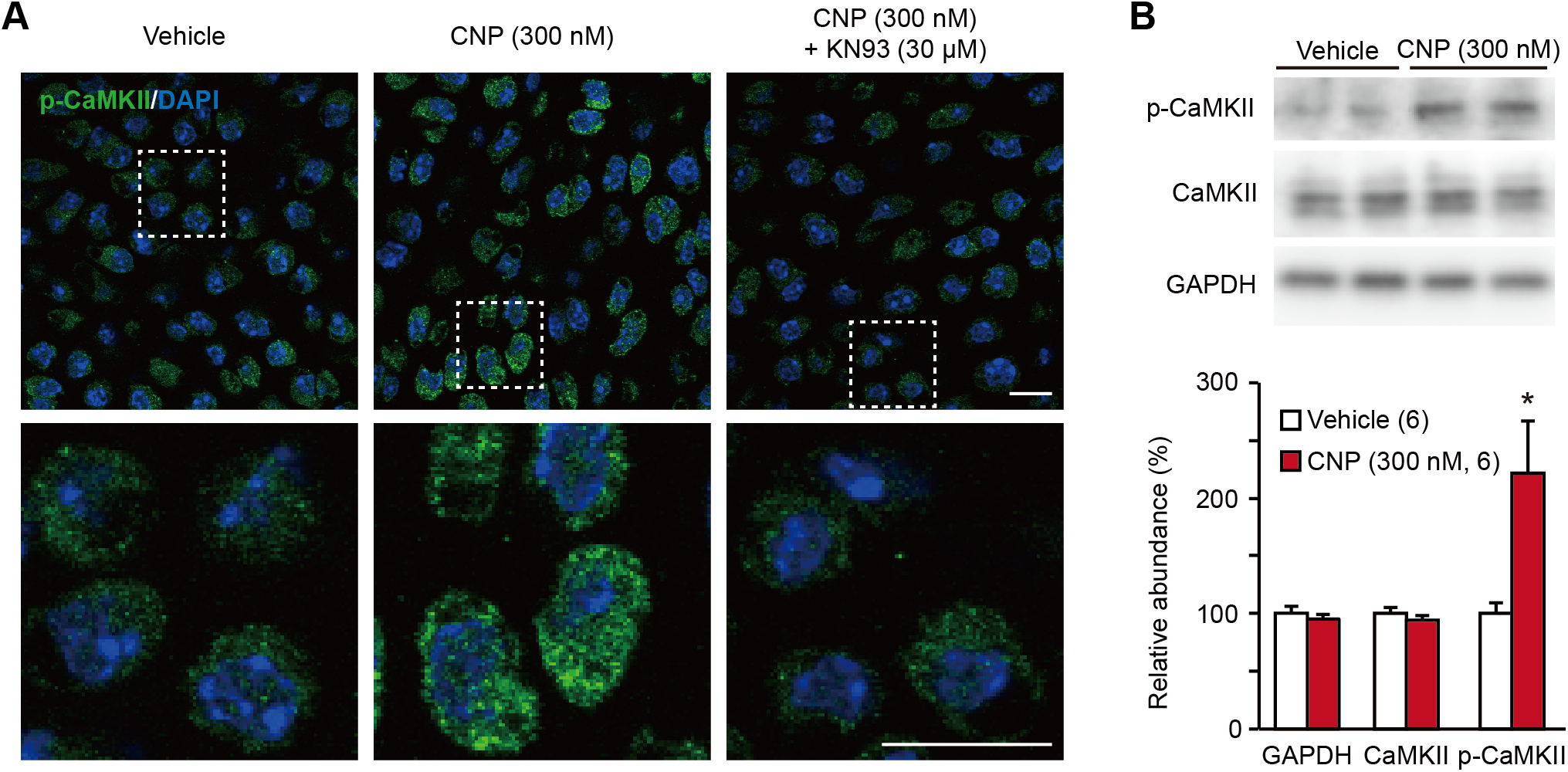
CaMKII activation in CNP-treated round chondrocytes. **(A)** Immunohistochemical staining against phospho-CaMKII (p-CaMKII) in round chondrocytes. Wild-type bone slices were pretreated with or without CNP and the CaMKII inhibitor KN93, and then subjected to immunostaining with antibody to p-CaMKII. DAPI (4’, 6-diamidino-2-phenylindole) was used for nuclear staining. Lower panels show high-magnification views of white-dotted regions in upper panels (scale bars, 10 μm). **(B)** Immunoblot analysis of total CaMKII and p-CaMKII in growth plate cartilage. Growth plate lysates were prepared from wild-type bone slices pretreated with or without CNP, and subjected to immunoblot analysis with antibodies against total CaMKII and p-CaMKII (upper panel). Glyceraldehyde-3-phosphate dehydrogenase (GAPDH) was also analyzed as a loading control. The immunoreactivities observed were densitometrically quantified and are summarized (lower graph). A significant difference between CNP- and vehicle-pretreatments is marked with an asterisk (\**p*<0.05 in one-way ANOVA and Tukey’s test). The data are presented as the means ± SEM. with *n* values indicating the number of examined mice.

### Pharmacologically activated BK channels facilitate bone outgrowth

Based on the present data from *in vitro* experiments, the novel CNP-signaling route, represented as the NPR2-PKG-BK channel-TRPM7 channel-CaMKII axis, can be proposed in growth plate chondrocytes. We attempted to examine the proposed signaling axis in metatarsal bone culture, a widely used *ex vivo* model system for analyzing bone growth and endochondral ossification [24]. In chondrocyte-specific *Trpm7*-knockout mice (*Trpm7*^fl/fl^, *11Enh*-Cre^+/-^), Cre recombinase is expressed under the control of the collagen type XI gene enhancer and promoter, and thus inactivates the floxed *Trpm7* alleles in cartilage cells [11]. The bone rudiments prepared from control embryos (*Trpm7*^fl/fl^, *11Enh-Cre*^-/-^) regularly elongated during *ex vivo* culture, and their outgrowth was significantly stimulated by the supplementation with CNP (30 nM) into the culture medium (Figure 7A). In contrast, the mutant rudiments prepared from the chondrocyte-specific *Trpm7*-knockout embryos were reduced in initial size and did not respond to the CNP supplementation. Therefore, CNP-facilitated bone outgrowth seems to require TRPM7 channels expressed in growth plate chondrocytes.

**Figure 7.**
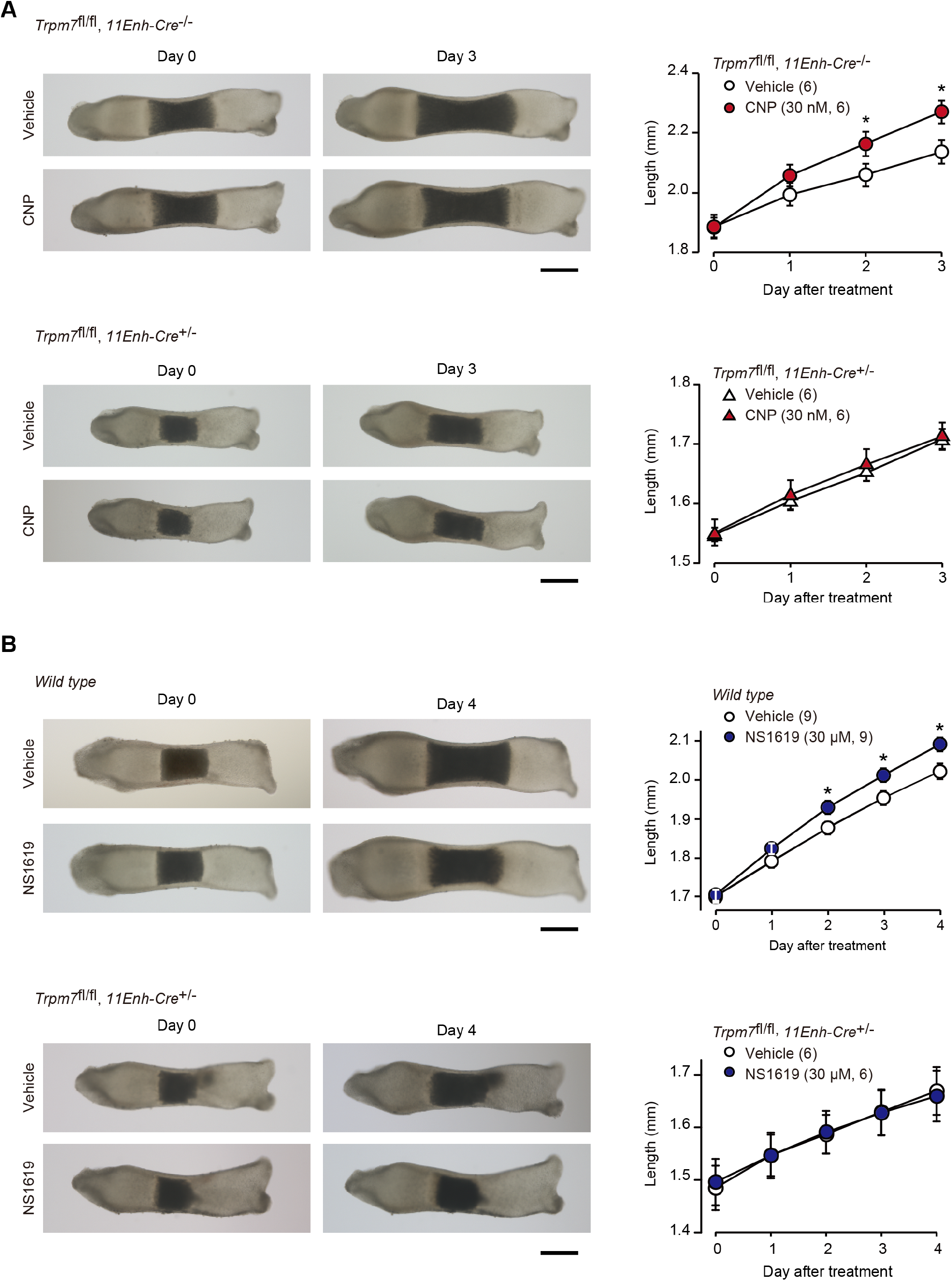
Contribution of TRPM7 and BK channels to CNP-facilitated bone outgrowth. (**A**) Loss of CNP-facilitated outgrowth in *Trpm7*-deficient bones. Metatarsal rudiments isolated from the chondrocyte-specific *Trpm7*-knockout (*Trpm7*^fl/fl^, *11Enh-Cre*^+/-^) and control (*Trpm7*^fl/fl^, *11Enh-Cre*^-/-^) embryos were precultured in normal medium for 6 days, and then cultured in medium supplemented with or without CNP for 3 days. Representative images of cultured metatarsals are shown (left panels; scale bar, 0.3 mm), and longitudinal bone outgrowth during the CNP-supplemented period was statistically analyzed in each genotype group (right graphs). Significant CNP-supplemented effects are marked with asterisks (\**p*<0.05 in *t*-test). The data are presented as the means ± SEM. with *n* values indicating the number of examined mice. (**B**) Stimulated bone outgrowth by the BK channel activator NS1619. Metatarsal rudiments isolated from wild-type and the chondrocyte-specific *Trpm7*-knockout embryos were precultured in normal medium for 5 days, and then cultured in medium supplemented with or without NS1619 for 4 days. Representative images of cultured metatarsals are shown (left panels; scale bar, 0.3 mm), and longitudinal bone outgrowth during the NS1619-supplemented period was statistically analyzed in each genotype group (right graphs). A significant NS1619-supplemented effect is marked with asterisks (\**p*<0.05 in *t*-test). The data are presented as the means ± SEM. with *n* values indicating the number of examined mice.

In our proposed signaling axis, activated BK channels exert an essential role by converting the chemical signal into the electrical signal. We finally examined the effect of the BK channel activator NS1619 on bone outgrowth (Figure 7B). NS1619 supplementation (30 μM) significantly stimulated the outgrowth of wild-type bone rudiments. In contrast, under the same culture conditions, no stimulation was detected in the mutant rudiments from the chondrocyte-specific *Trpm7*-deficient embryos. The observations in the bone culture support our conclusion that CNP activates BK channels and thus facilitates TRPM7-mediated Ca^2+^ influx in growth plate chondrocytes, stimulating bone growth.

## Discussion

We reported that in growth plate chondrocytes, PLC and BK channels maintain autonomic TRPM7-mediated Ca^2+^ fluctuations, which potentiate chondrogenesis and bone growth by activating CaMKII [11]. Based on the present data, together with the previous reports, we proposed a new CNP signaling axis in growth plate chondrocytes (Figure 7-figure supplement 5A). CNP-induced NPR2 activation elevates cellular cGMP content and thus activates PKG, leading to the phosphorylation of BK channels. The resulting BK channel activation likely induces cellular hyperpolarization to facilitate TRPM7-mediated Ca^2+^ entry by enhancing the Ca^2+^ driving force, leading to CaMKII activation. Therefore, it is likely that CaMKII activity is physiologically regulated by BK channels as a key player of the CNP signaling cascade. In a recent genetic study, several patients carrying loss-of-function mutations in the *KCNMA1* gene encoding BK channel α subunit were characterized by a novel syndromic growth deficiency associated with severe developmental delay, cardiac malformation, bone dysplasia and dysmorphic features [25]. In the *KCNMA1*-mutated disorder, CNP signaling likely fails to facilitate TRPM7-mediated Ca^2+^ fluctuations in growth plate chondrocytes and resulting insufficient Ca^2+^ entry may lead to systemic bone dysplasia associated with stunted growth plate cartilage. On the other hand, the origin of CNP may still be ambiguous in the signaling scheme. Transgenic mice overexpressing CNP in a chondrocyte-specific manner develop a prominent skeletal overgrowth phenotype, suggesting autocrine CNP signaling [26]. However, several genechip data in public databases indicate that prepro-CNP mRNA is abundantly expressed in the placenta among embryonic tissues (for example, see the records under accession number GSE28277 in NCBI database). Therefore, it may be important to further examine which cell type primarily produces CNP to facilitate bone growth during embryonic development.

From a physiological point of view, it is interesting to note that the proposed CNP signaling axis has clear overlap with the nitric oxide (NO) and ANP/BNP signaling cascades for vascular relaxation [27–29]. In blood vessels, NO is produced by endothelial cells in response to various stimuli including shear stress and acetylcholine, and activates soluble guanylate cyclase in neighboring vascular smooth muscle cells. ANP and BNP are released from the heart in response to pathological stresses, such as atrial distension and pressure overload, and are delivered to activate the receptor guanylate cyclase NPR1 in vascular muscle. In either case, the resulting cGMP elevation followed by PKG activation induces BK channel-mediated hyperpolarization and thus inhibits L-type Ca^2+^ channel gating, leading to vascular dilation due to decreased Ca^2+^ entry into vascular muscle. Therefore, activated BK channels inhibit the voltage-dependent Ca^2+^ influx in vascular muscle cells regarded as excitable cells (Figure 7-figure supplement 5B). In contrast, activated BK channels reversely stimulate TRPM7-mediated Ca^2+^ entry in growth plate chondrocytes classified as nonexcitable cells, because the channel activity is voltage-independently maintained by the intrinsic PI turnover rate.

CNP is an effective therapeutic reagent for achondroplasia and divergent short statures [26, 30, 31], and the phase III clinical trial of CNP therapy is now ongoing [2]. The proteins contributing to the CNP signaling axis may be new pharmaceutical targets for developing medications; in addition to NPR2, BK and TRPM7 channels are reasonably considered promising targets. Moreover, phosphodiesterase subtypes might be useful targets, although the subtypes responsible for cGMP hydrolysis remain to be identified in growth plate chondrocytes. Chemical compounds specifically targeting the signaling axis defined in this study would be useful drugs for not only clinical treatment of developmental disorders but also artificially modifying body sizes in farm and pet animals.

## Materials and Methods

### Reagents, primers and mice

Reagents and antibodies used in this study are listed in Key Resources Table. Synthetic primers used for RT-PCR analysis and mouse genotyping are also listed in Key Resourses Table. C57BL mice were used as wild-type mice in this study. Chondrocyte-specific *Trpm7-*knockout mice with C57BL genetic background were generated and genotyped as previously described [11]. Chondrocyte-specific *Npr2*-knockout mice with C57BL background were generated as previously described [12], and we newly designed primers for detecting the *Col2a1-Cre* transgene and the floxed *Npr2* gene in this study (Figure 1-figure supplement 1). All experiments in this study were conducted with the approval of the Animal Research Committee according to the regulations on animal experimentation at Kyoto University.

### Bone slice preparations

Femoral bones were isolated from E17.5 mice and immersed in a physiological salt solution (PSS): (in mM) 150 NaCl, 4 KCl, 1 MgCl_2_, 2 CaCl_2_, 5.6 glucose, and 5 HEPES (pH 7.4). Longitudinal bone slices (∼40 µm thickness) were prepared using a vibrating microslicer (DTK-1000N, Dosaka EM Co., Japan) as previously described [11].

### Ca^2+^ imaging

Fura-2 Ca^2+^ imaging of bone slices was performed as previously described [11]. Briefly, bone slices placed on glass-bottom dishes (Matsunami, Japan) were incubated in PSS containing 15 μM Fura-2AM for 1 hr at 37°C. For ratiometric imaging, excitation light of 340 and 380 nm was alternately delivered, and emission light of >510 nm was detected by a cooled EM-CCD camera (Model C9100-13; Hamamatsu Photonics, Japan) mounted on an upright fluorescence microscope (DM6 FS, Leica, Germany) using a 40x water-immersion objective (HCX APO L, Leica). In typical measurements, ∼30 round chondrocytes were randomly examined in each slice preparation to select the Ca^2+^ fluctuation-positive cells generating spontaneous events (>0.025 in Fura-2 ratio) using commercial software (Leica Application Suite X), and recording traces from the positive cells were then analyzed using Fiji/ImageJ software (US. NIH) for examining Ca^2+^ fluctuation amplitude and frequency. Imaging experiments were performed at room temperature (23-25 °C) and PSS was used as the normal bathing solution. For the pretreatments of CNP, ANP and 8-pCPT-cGMP, bone slices were immersed in PSS with the indicated compound for 1 hr at room temperature after Fura-2 loading.

### Membrane potential monitoring

Bone slices were perfused with the PSS containing 200 nM oxonol VI at room temperature and analyzed as previously described [33]. To prepare the calibration plot showing the relationship between the fluorescence intensity and membrane potential, saline solutions containing 20 mM, 40 mM, 60 mM or 100 mM KCl were used as bathing solutions. Fluorescence images with excitation at 559 nm and emission at >606 nm were captured at a sampling rate of ∼7.0 s using a confocal laser scanning microscope (FV1000; Olympus).

### Immunochemical analysis of CaMKII

Bone slices were pretreated with or without CNP were subjected to immunochemical assessments as previously described [34]. Briefly, for immunohistochemical analysis, bone slices were fixed in 4% paraformaldehyde and treated with 1% hyaluronidase to enhance immunodetection [35, 36]. After blocking with fetal bovine serum-containing solution, bone slices were reacted with primary and Alexa 488-conjugated secondary antibodies and observed with a confocal microscope (FV1000; Olympus). For immunoblot analysis, bone slices were lysed in the buffer containing 4% sodium deoxycholate, 20 mM Tris-HCl (pH 8.8) and a phosphatase inhibitor cocktail (100 mM NaF, 10 mM Na_3_PO_4_, 1 mM Na_2_VO_3_ and 20 mM β-glycerophosphate). The resulting lysate proteins were electrophoresed on SDS-polyacrylamide gels and electroblotted onto nylon membranes for immunodetection using primary and HRP-conjugated secondary antibodies. Antigen proteins were visualized using a chemiluminescence reagent and image analyzer (Amersham Imager 600, Cytiva). The immunoreactivities yielded were quantitatively analyzed by means of Fiji/ImageJ software.

### Metatarsal organ culture

Metatarsal bone rudiments were cultured as previously described [24]. Briefly, the three central metatarsal rudiments were dissected from E15.5 mice and cultured in αMEM containing 5 µg/ml ascorbic acid, 1 mM β-glycerophosphate pentahydrate, 100 units/ml penicillin, 100 µg/ml streptomycin and 0.2% bovine serum albumin (fatty acid free). The explants were analyzed under a photomicroscope (BZ-X710, Keyence, Japan) for size measurements using Fiji/ImageJ software.

### Gene expression analysis

Quantitative RT-PCR analysis was performed as previously described [37]. Total RNA was prepared from mouse tissues using a commercial reagent (Isogen) and reverse-transcribed using a commercial kit (ReverTra ACE qPCR-RT kit). The resulting cDNAs were examined by real-time PCR (LightCycler 480 II, Roche), and the cycle threshold was determined from the amplification curve as an index for relative mRNA content in each reaction.

### Quantification and statistical analysis

All data obtained are presented as the means ± SEM. with *n* values indicating the number of examined mice. Student *t*-test and ANOVA were used for two-group and multiple group comparisons, respectively (Prism 7, GraphPad Software Inc.): *p*<0.05 was considered to be statistically significant.

## Acknowledgements

We thank Jun Matsushita (Graduate School of Pharmaceutical Sciences, Kyoto University) for mouse *in vitro* fertilization. This work was supported in part by the MEXT/JSPS (KAKENHI Grant Number 21H02663 and 20H03802), Platform Project for Supporting Drug Discovery and Life Science Research (JP19am0101092j0003), Takeda Science Foundation, Kobayashi International Scholarship Foundation, the NAKATOMI Foundation, Vehicle Racing Commemorative Foundation and Japan Foundation for Applied Enzymology.

## Author contributions

Y.M. and A.I. are equally contributing first authors. Y.M., A.I. and N.O. conducted Ca^2+^ imaging analysis. Y.M., A.I., R.K., T.Y., Y.U., I.Y., S.K. and M.N. conducted biochemical and cell physiological analysis. Y.M. and F.L. conducted membrane potential measurement. Y.U., I.Y., T.H., K.N., Y.M., H.A., and K.N. generated conditional knockout mice. R.K. and T.Y. conducted metatarsal organ culture analysis. Y.M., A.I. and H.T. drafted the manuscript. H.T. oversaw this project.

## Figure legends for figure supplements

**Figure 1-figure supplement 1.**
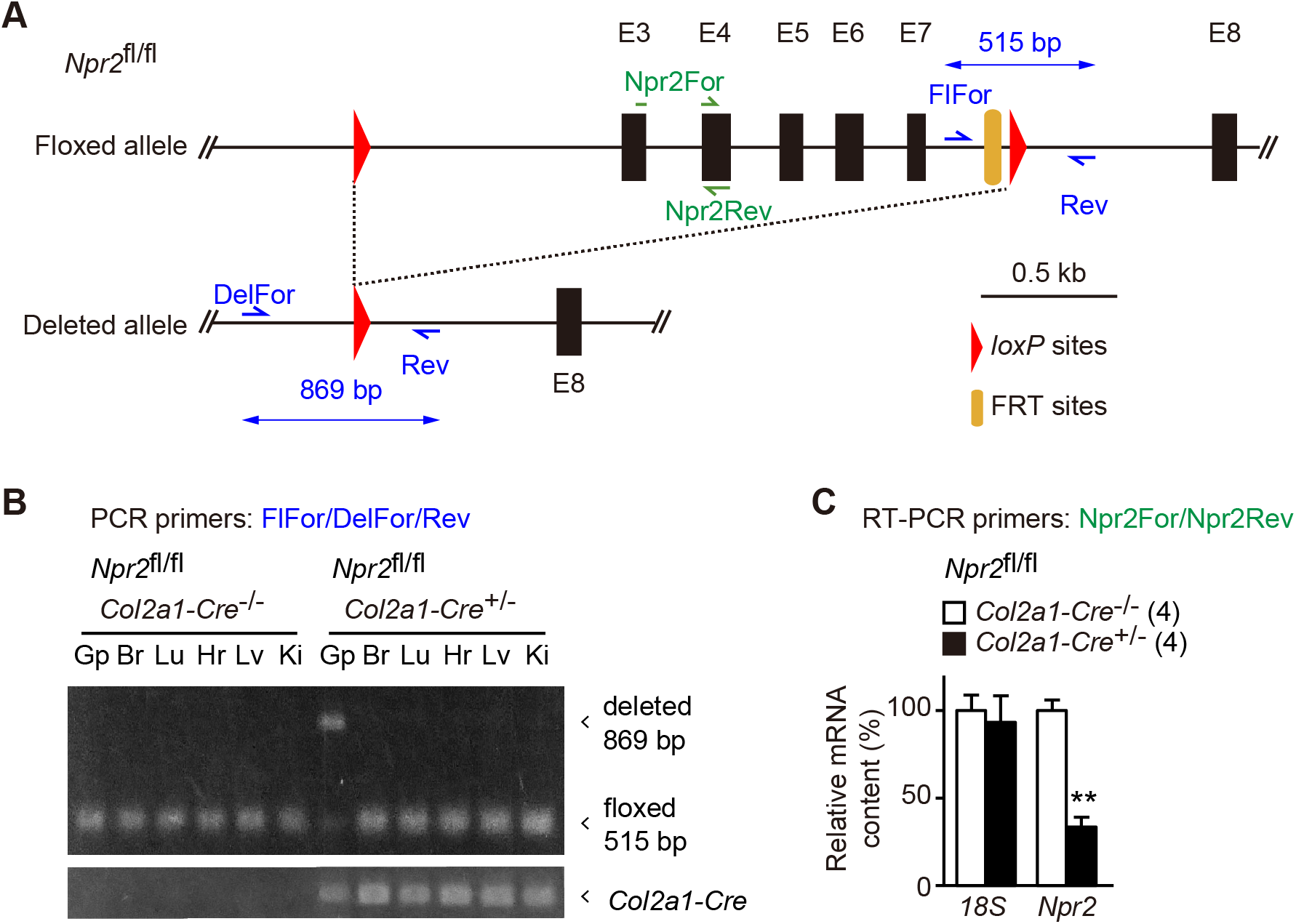
Chondrocyte-specific *Npr2 ablation*. **(A)** Organization of floxed and deleted *Npr2* alleles. The chondrocyte-specific *Npr2*-knockout (*Npr2*^fl/fl^, *Col2a1-Cre*^+/-^) mice were previously generated [12]. In this study, genotyping primers were newly designed, and *Npr2* ablation was evaluated in growth plates. The genomic map shows PCR primers for detecting the mutated *Npr2* alleles and *Npr2* mRNA. **(B)** *Npr2* gene ablation in various tissues from the chondrocyte-specific *Npr2*-knockout mice. Genomic DNAs were prepared from tissues (Gp, humeral growth plate; Br, brain; Lu, lung; Hr, heart; Lv, liver; Ki, kidney) from the E17.5 chondrocyte-specific *Npr2*-knockout and control embryos, and subjected to PCR analysis to detect the floxed and deleted *Npr2* alleles; the *Col2a1-Cre* transgene was also examined. **(C)** Reduction of *Npr2* mRNA in mutant growth plates prepared from the chondrocyte-specific *Npr2*-knockout mice. Total RNAs were prepared from humeral growth plates from the E17.5 embryos, and subjected to RT-PCR analysis for estimating *Npr2* mRNA content. 18S ribosomal RNA was examined as an internal control. The relative mRNA contents were estimated from cycle thresholds in RT-PCR reactions and are summarized in the bar-graph. The data represent means ± SEM, and the numbers of mice examined are shown in parentheses. A significant difference between the genotype is marked with an asterisk (\*\**p*<0.01 in *t*-test).

**Figure 1-figure supplement 2.**
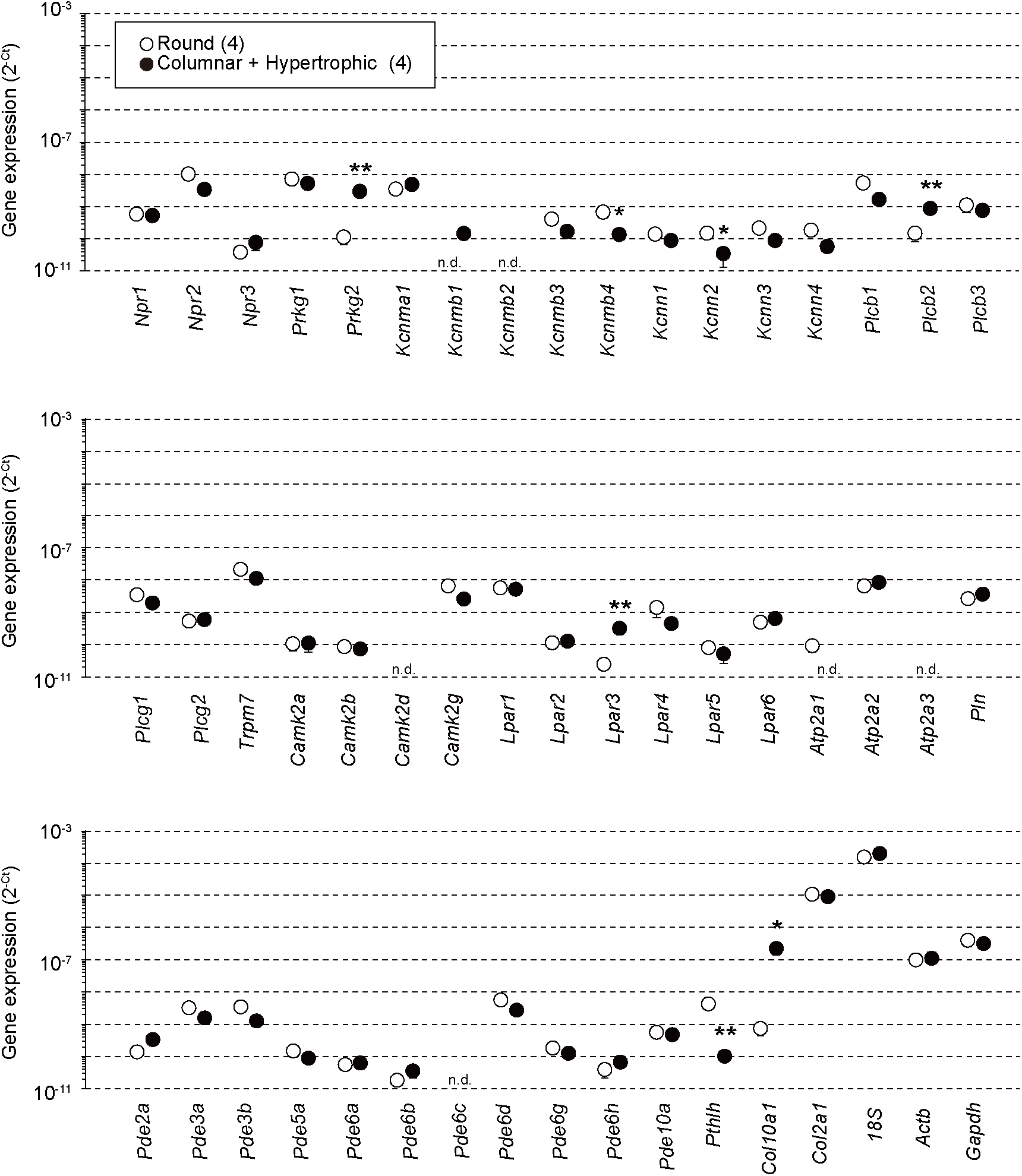
Gene expression analysis in wild-type growth plate chondrocytes. Total RNAs were prepared from growth plate sections packed with round chondrocytes or enriched with columnar and hypertrophic chondrocytes, and subjected to RT-PCR analysis. The cycle threshold (Ct) was determined for each RT-PCR reaction for estimating relative mRNA content. The data represent the mean ± SEM, and the numbers of mice examined are shown in parentheses. Significant differences between the growth plate sections are marked with asterisks (\**p*<0.05 and \*\**p*<0.01 in *t*-test). n.d.: not detectable.

**Figure 3-figure supplement 3.**
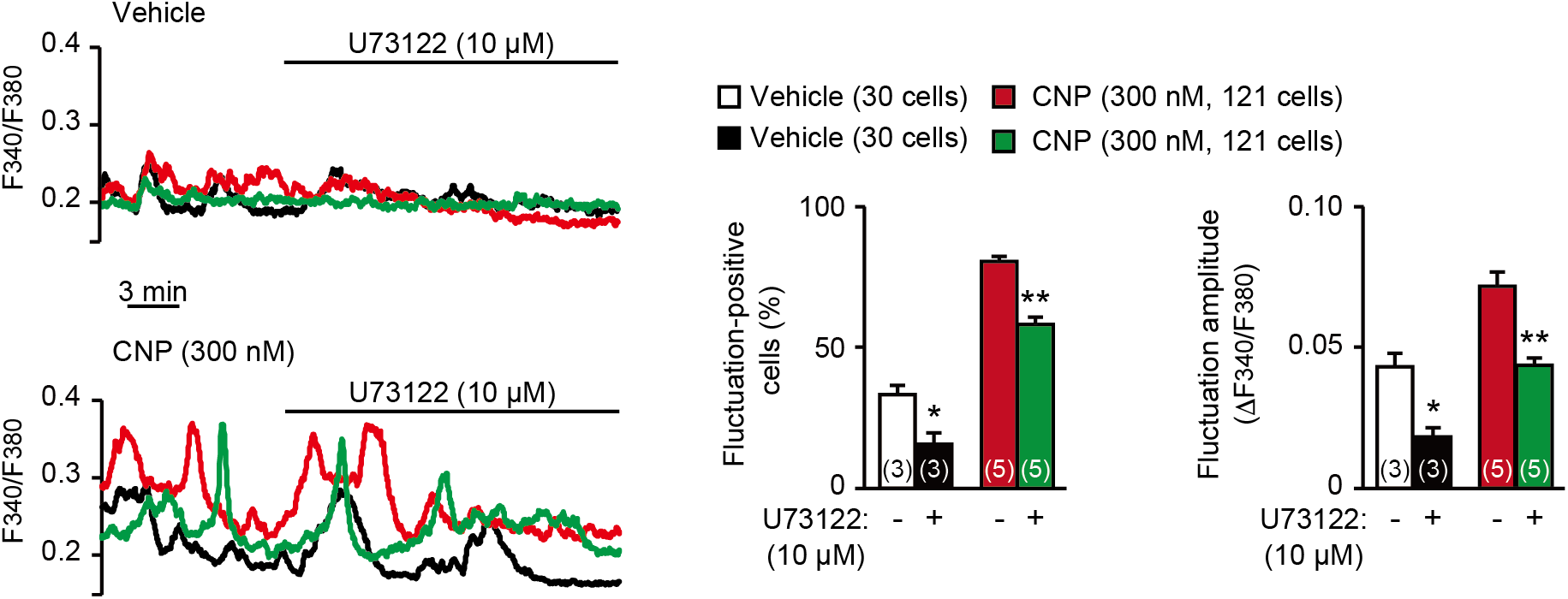
Effects of PLC inhibitor U73122 on CNP-facilitated Ca^2+^ fluctuations. In Ca^2+^ imaging, U73122 was bath-applied to wild-type round chondrocytes pretreated with or without CNP. Representative recording traces are shown (left panels), and the effects of U73122 are summarized (right bar-graphs). Data represent means ± SEM, and the numbers of cells and mice examined are shown in parentheses in the keys and graph bars, respectively. Significant differences between before and after the U73122 treatment are marked with asterisks (\**p*<0.05 and \*\**p*<0.01 in one-way ANOVA and Tukey’s test).

**Figure 3-figure supplement 4.**
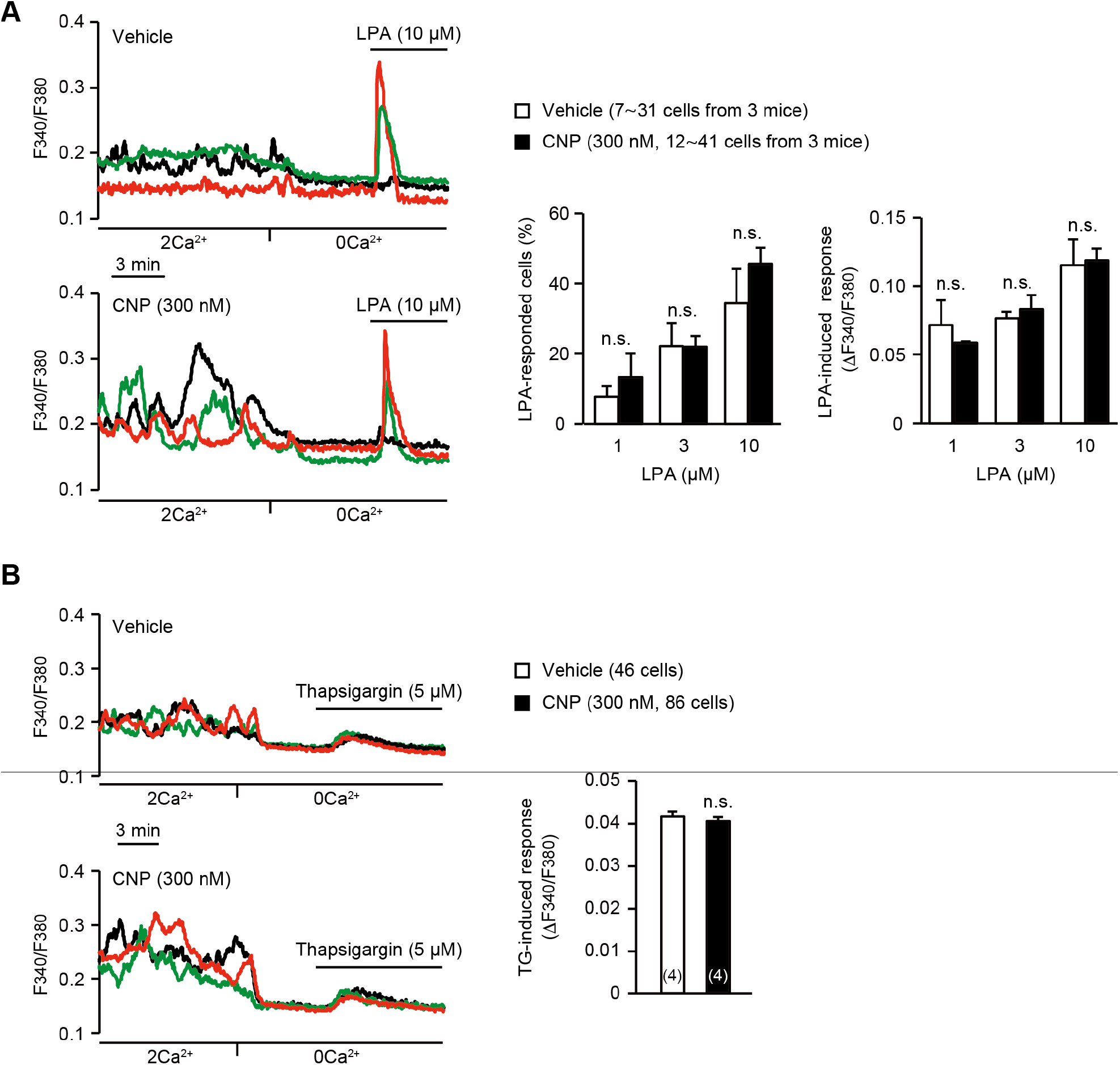
Store Ca^2+^ release in CNP-treated round chondrocytes. **(A)** Store Ca^2+^ release triggered by 1-oleoyl lysophosphatidic acid (LPA) in wild-type round chondrocytes pretreated with or without CNP. Representative recording traces are shown (left panels), and LPA-evoked Ca^2+^ responses are summarized (right graphs). Data represent means ± SEM, and the numbers of cells and mice examined are shown in parentheses in the keys and graph bars, respectively. No significant differences were observed between CNP- and vehicle-pretreated groups (one-way ANOVA and Tukey’s test). **(B)** Ca^2+^ leak responses evoked by the SERCA pump inhibitor thapsigargin (TG) in wild-type round chondrocytes pretreated with or without CNP. Representative recording traces are shown (left panels), and TG-evoked Ca^2+^ responses are summarized (right bar-graphs). Data represent means ± SEM, and the numbers of cells and mice examined are shown in parentheses in the keys and graph bars, respectively. No significant differences were observed between CNP- and vehicle-pretreated groups (*t*-test).

**Figure 7-figure supplement 5.**
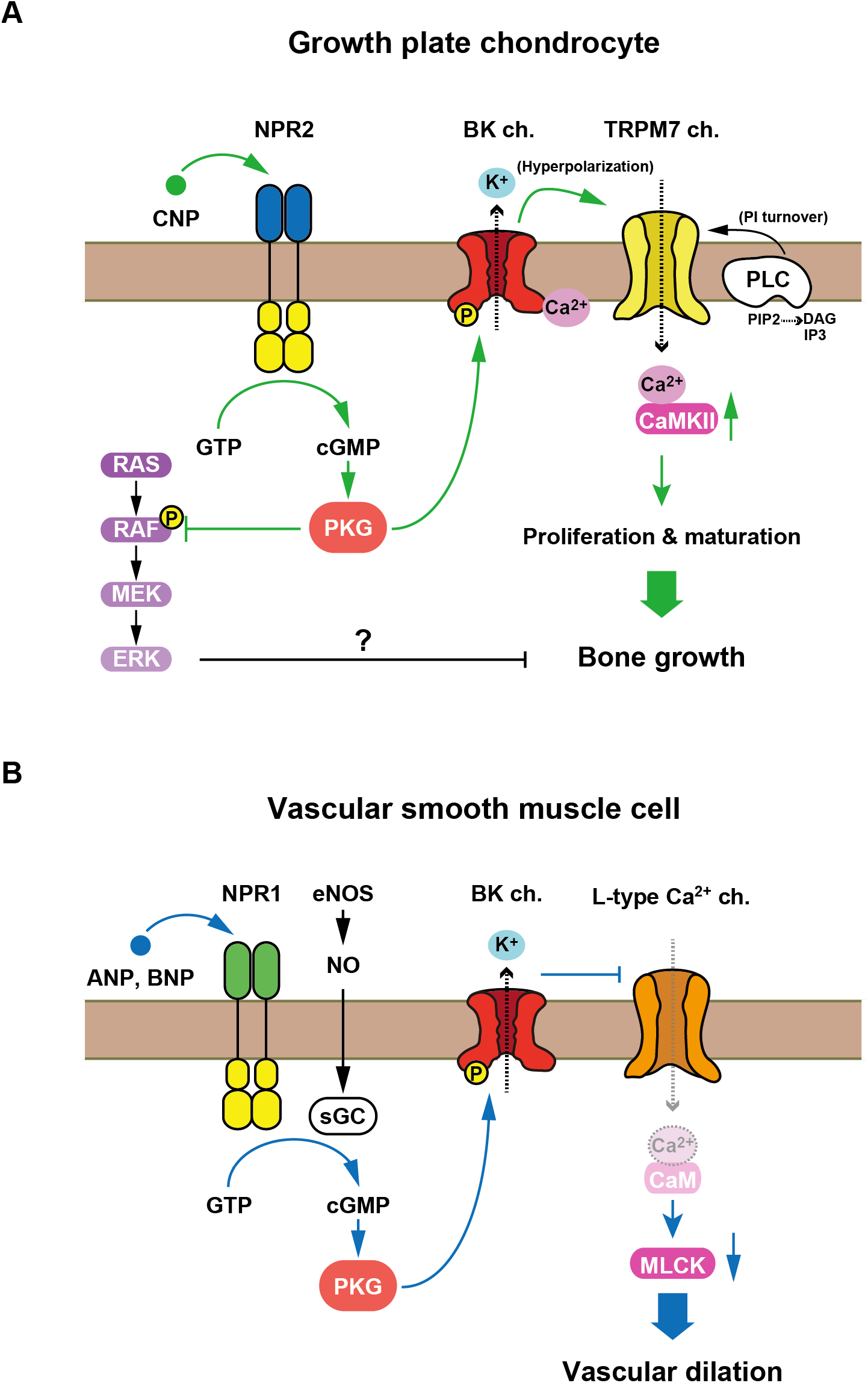
Proposed CNP-evoked signaling in growth plate chondrocytes. **(A)** The schematic diagram representing the NPR2-PKG-BK channel-TRPM7 channel-CaMKII axis proposed as an essential CNP signaling cascade in growth plate chondrocytes. Previous studies proposed that the RAF-MEK-ERK axis is also involved in growth plate CNP signaling [8]. **(B)** The schematic diagram representing the NO- and ANP/BNP-induced relaxation signaling in vascular smooth muscle.

## Key Resources Table

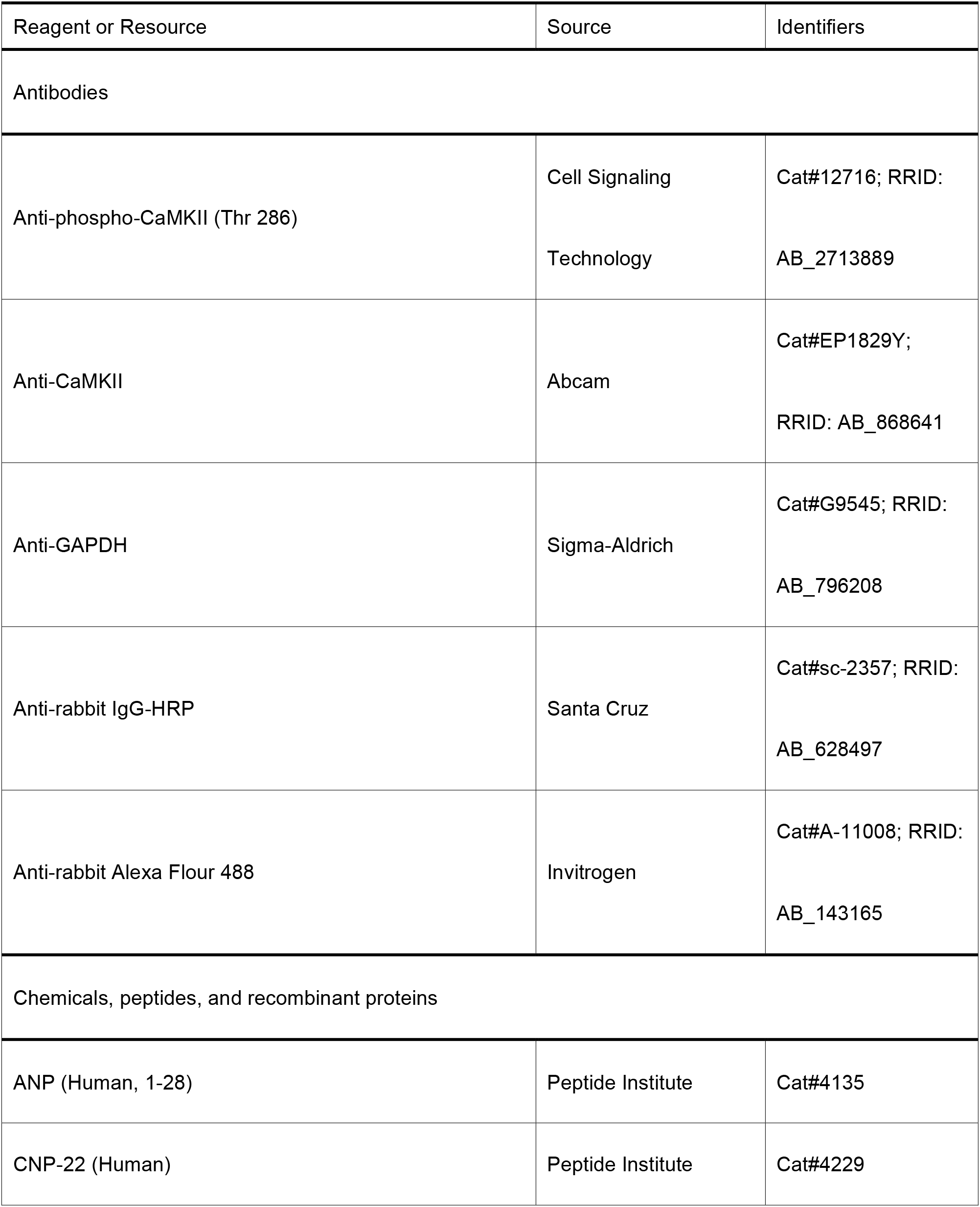
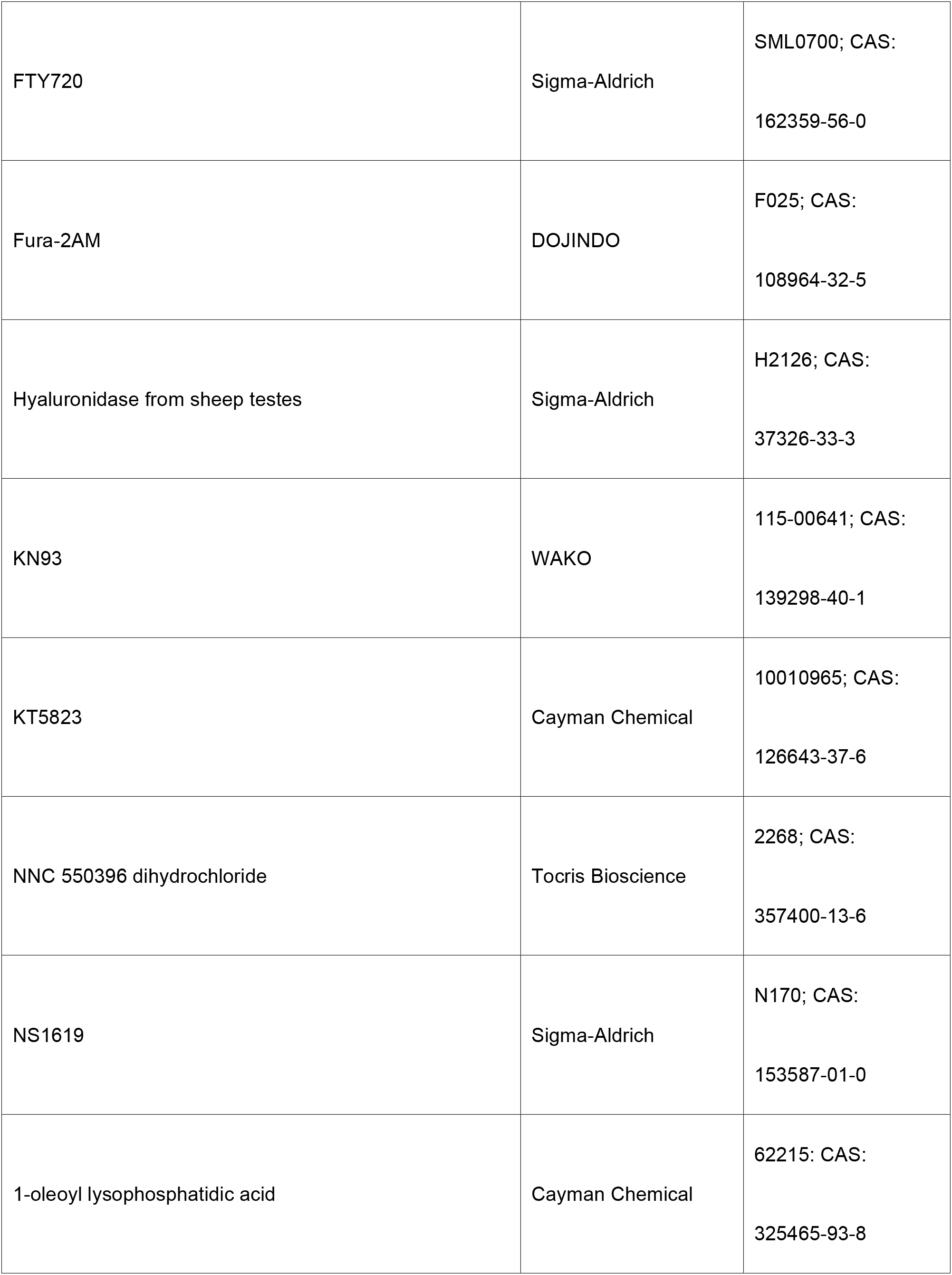
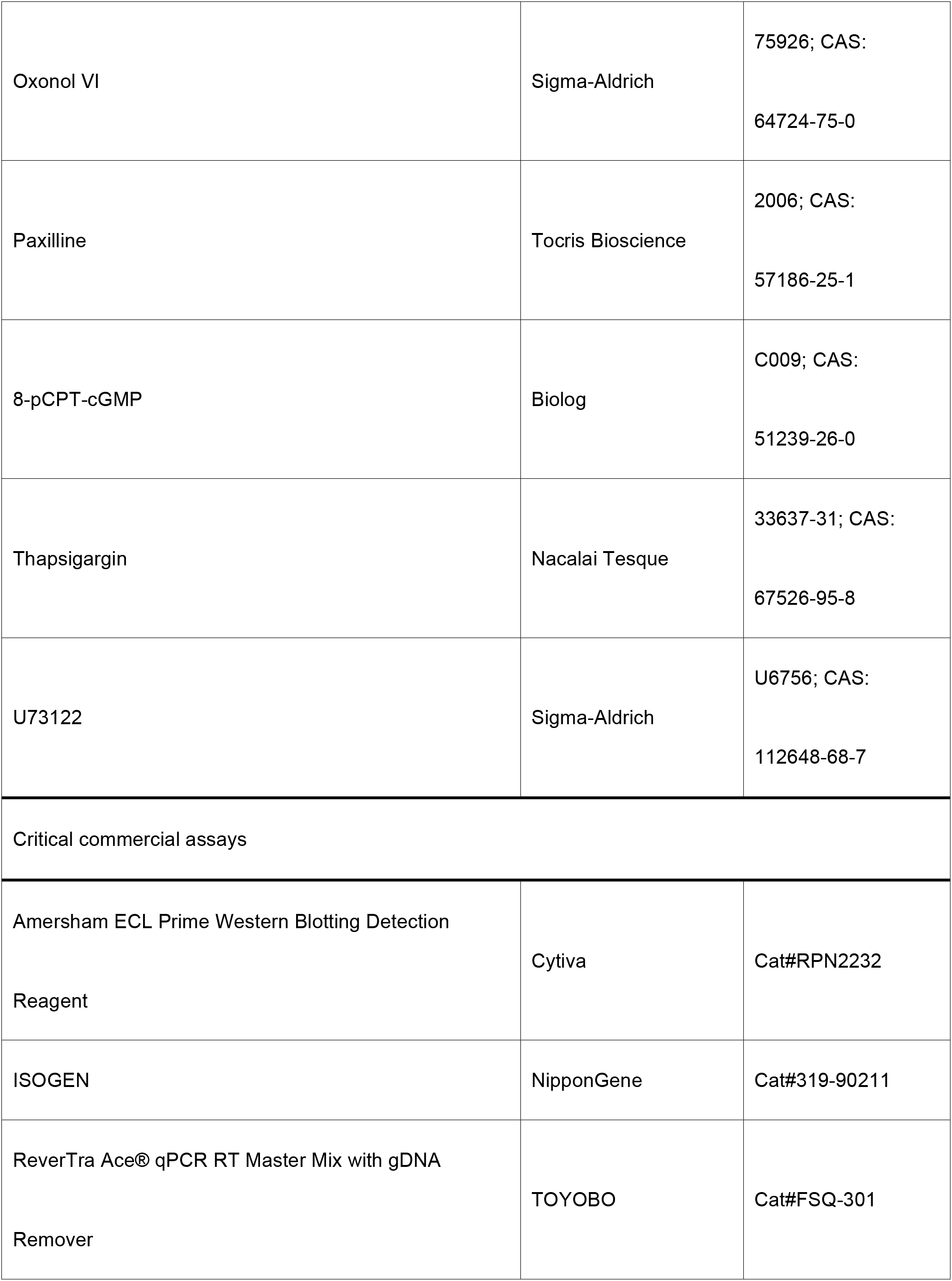
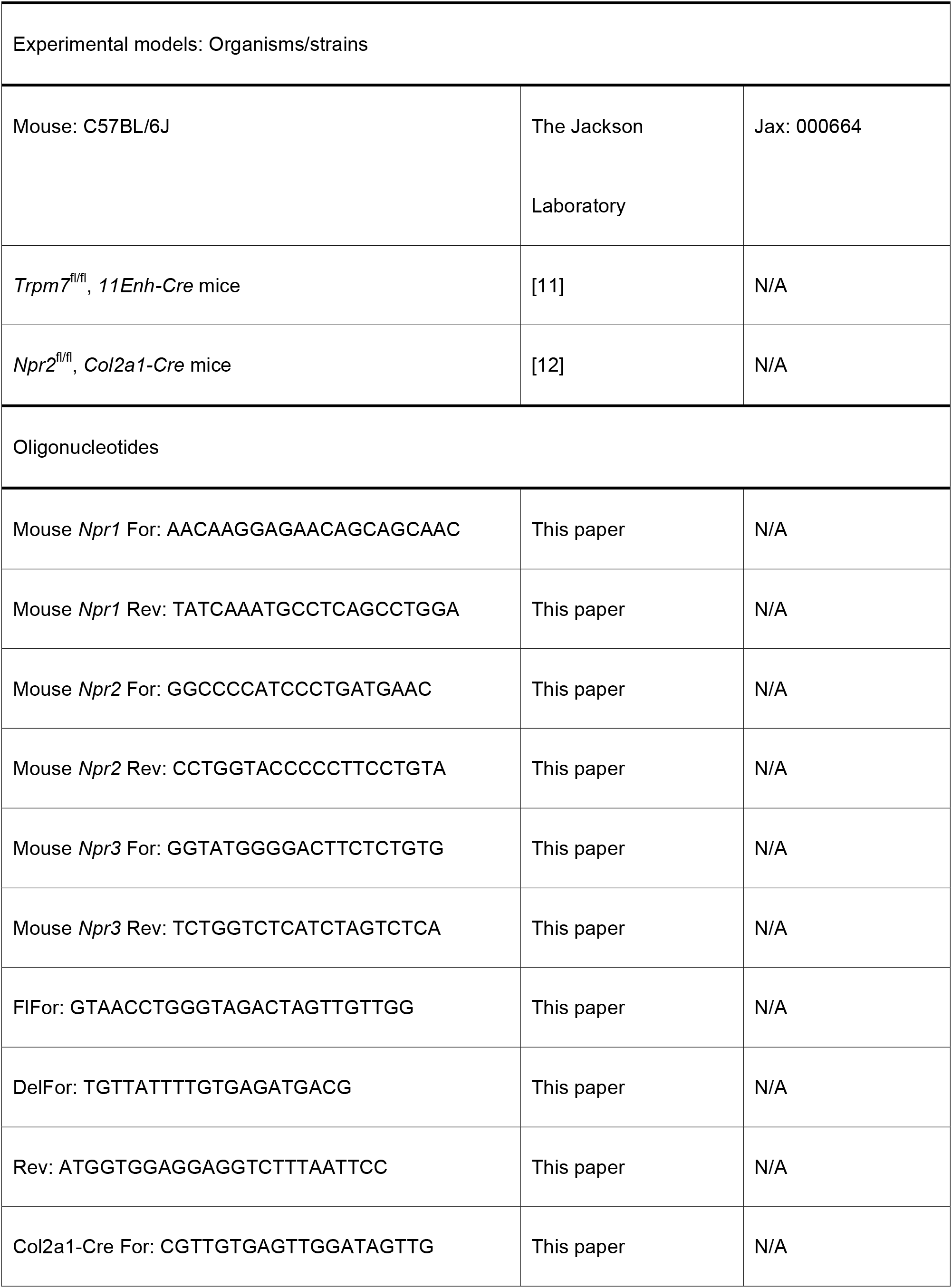
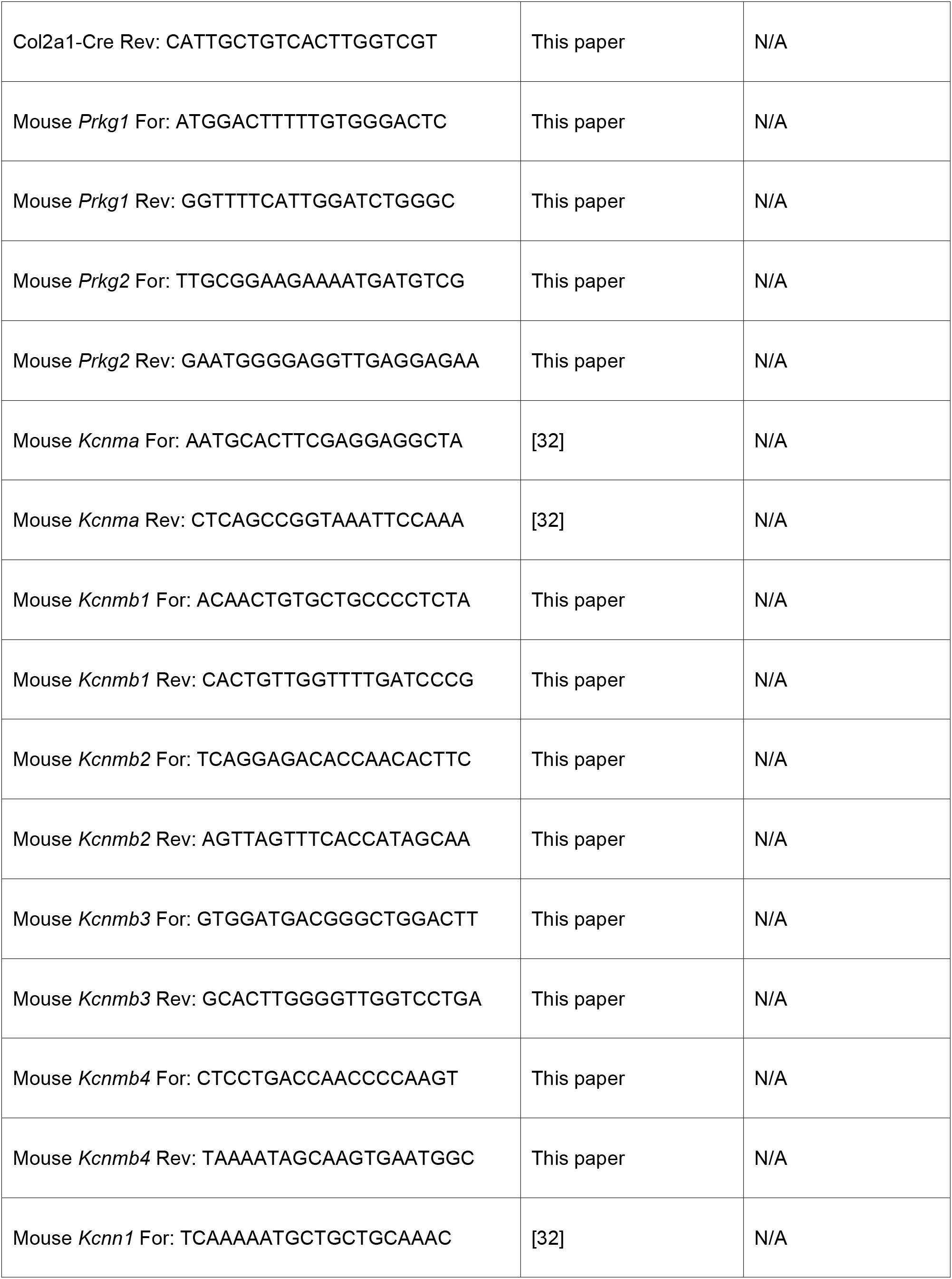
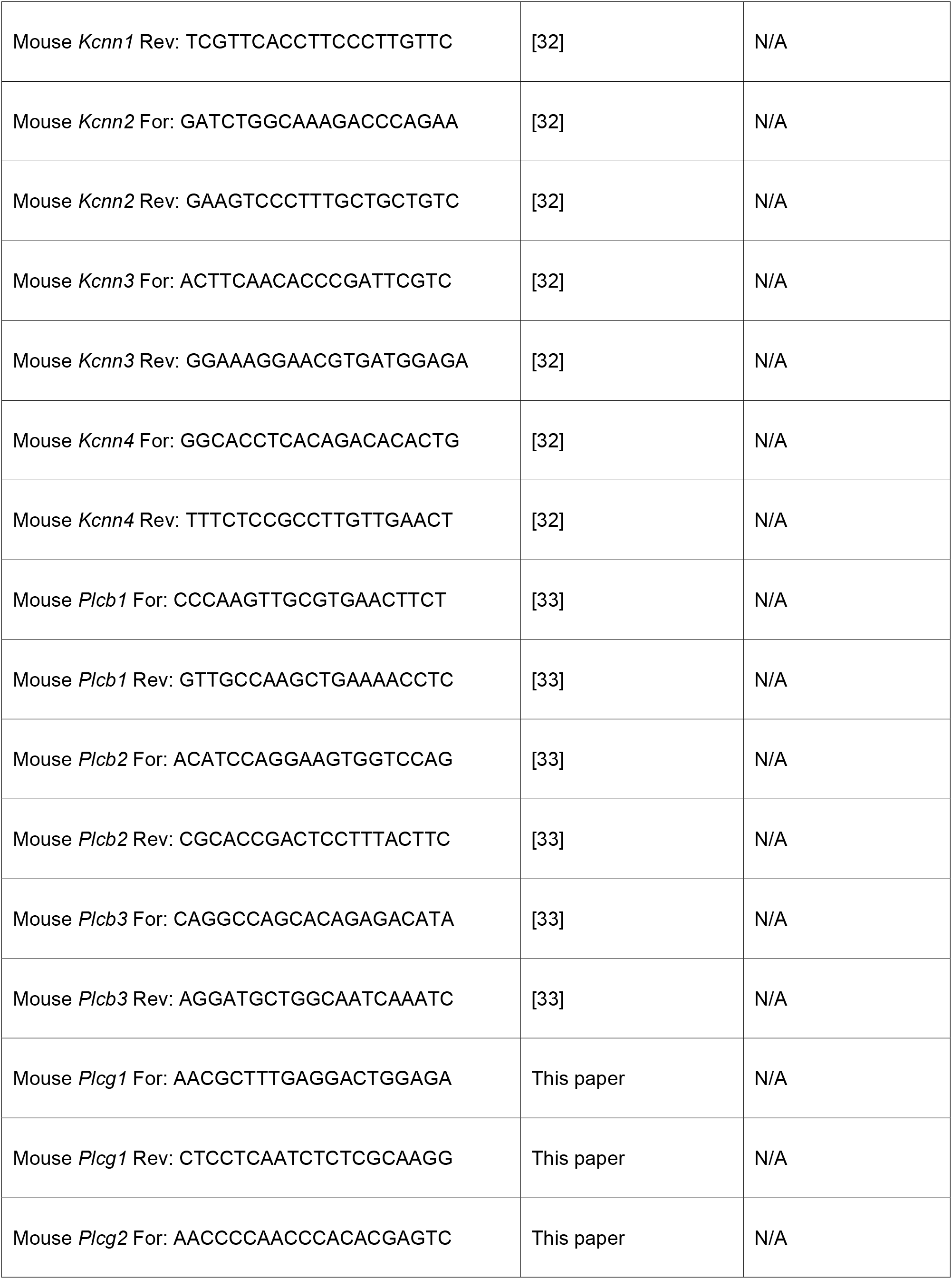
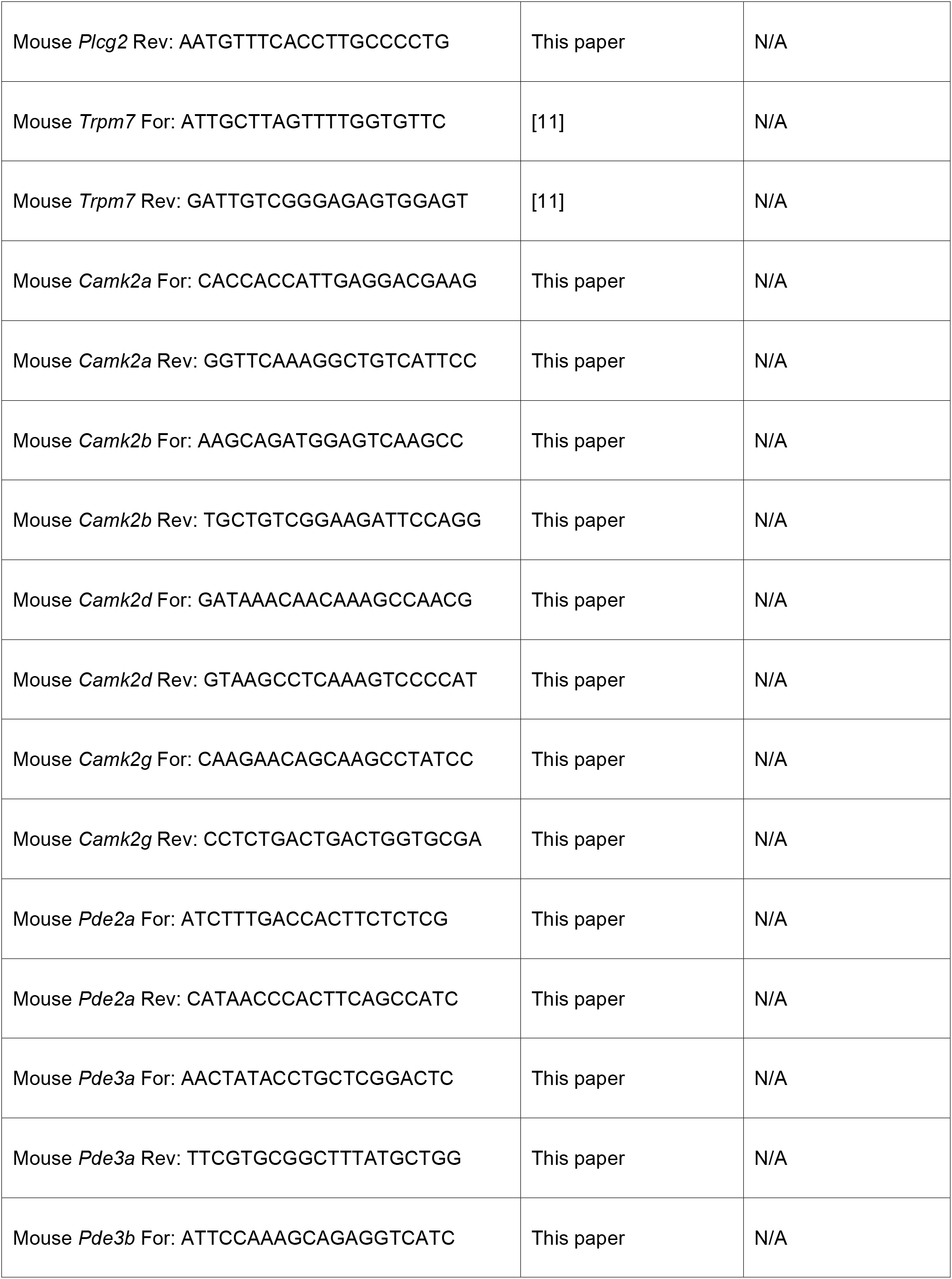
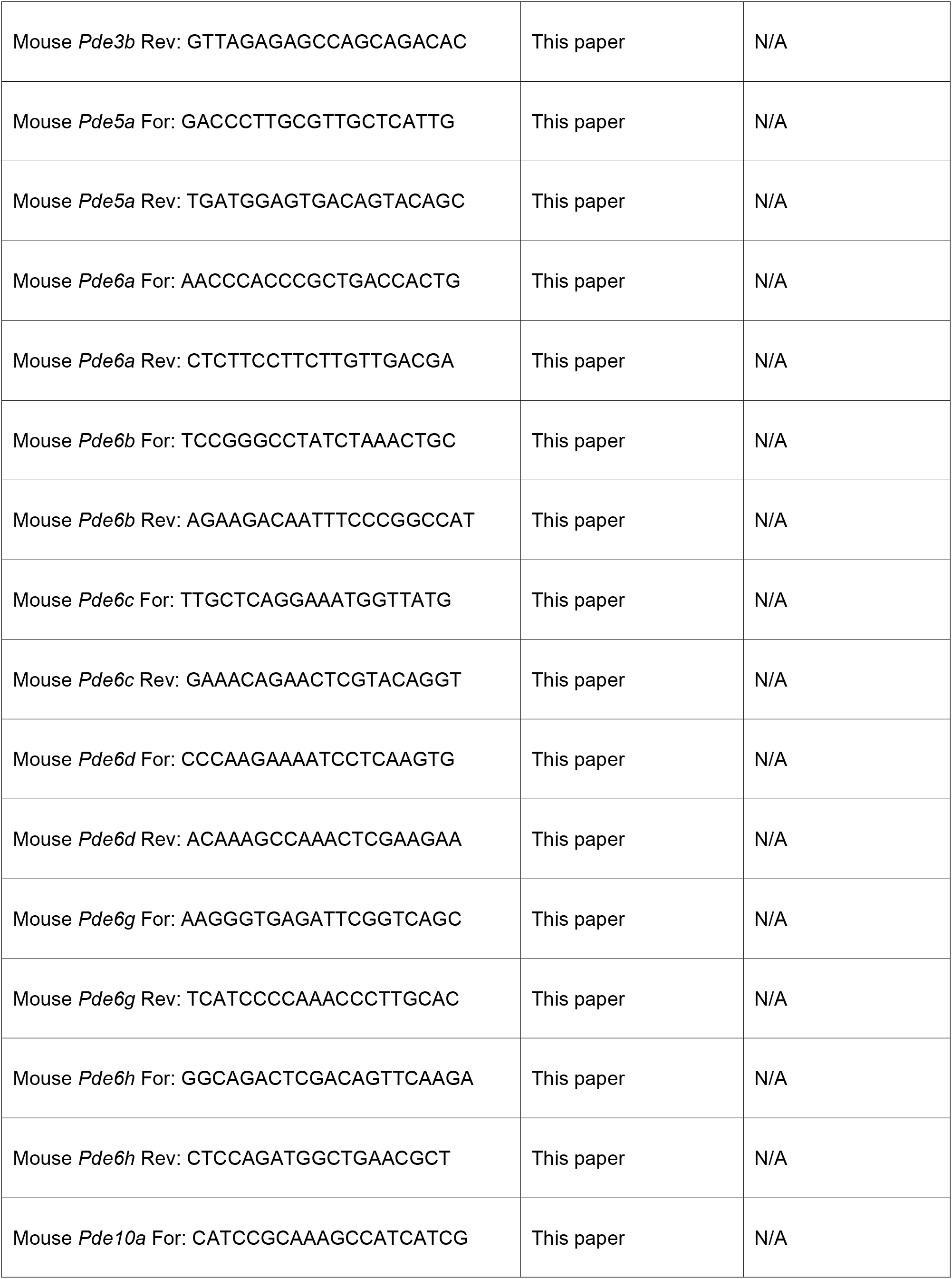
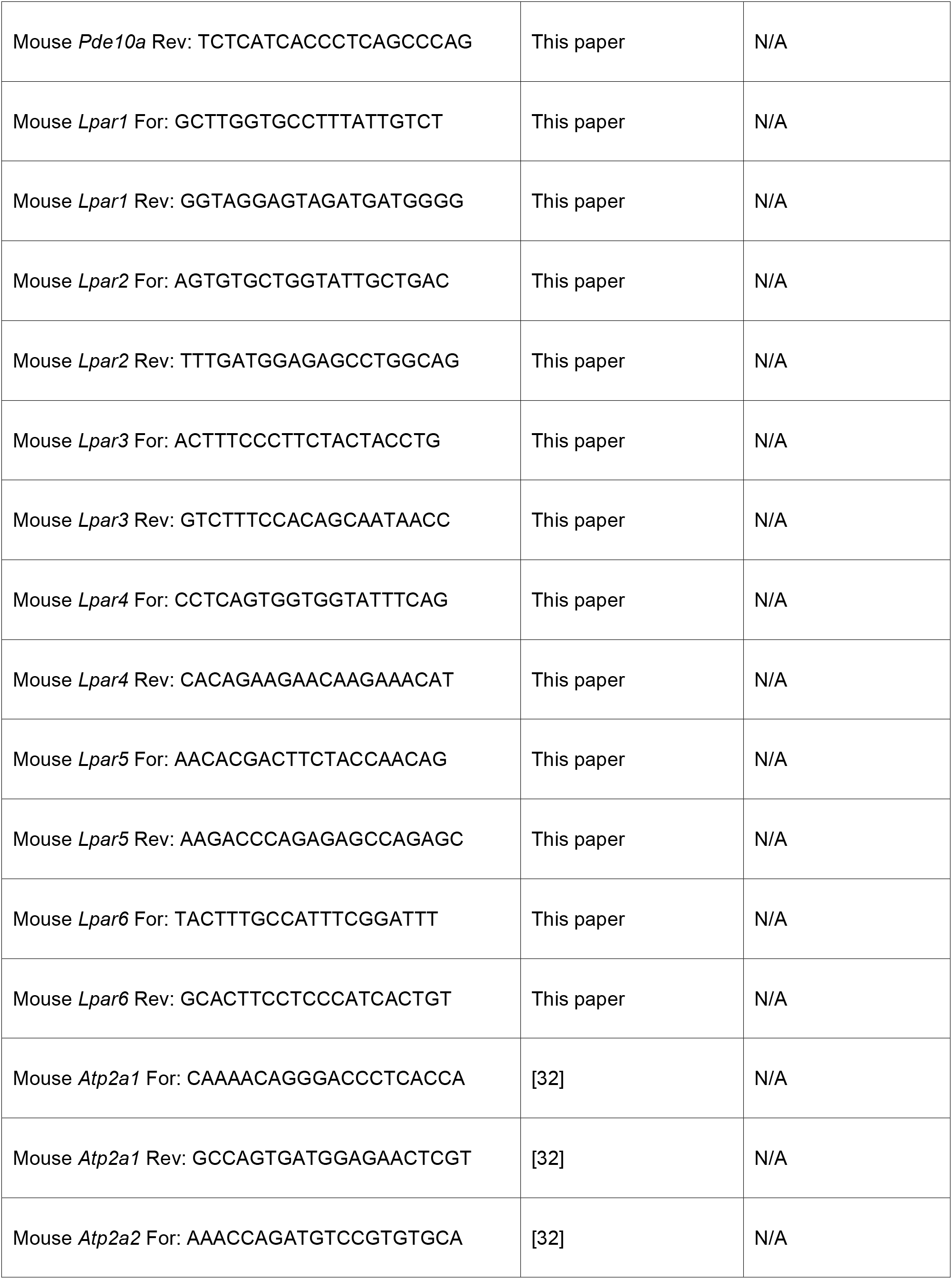
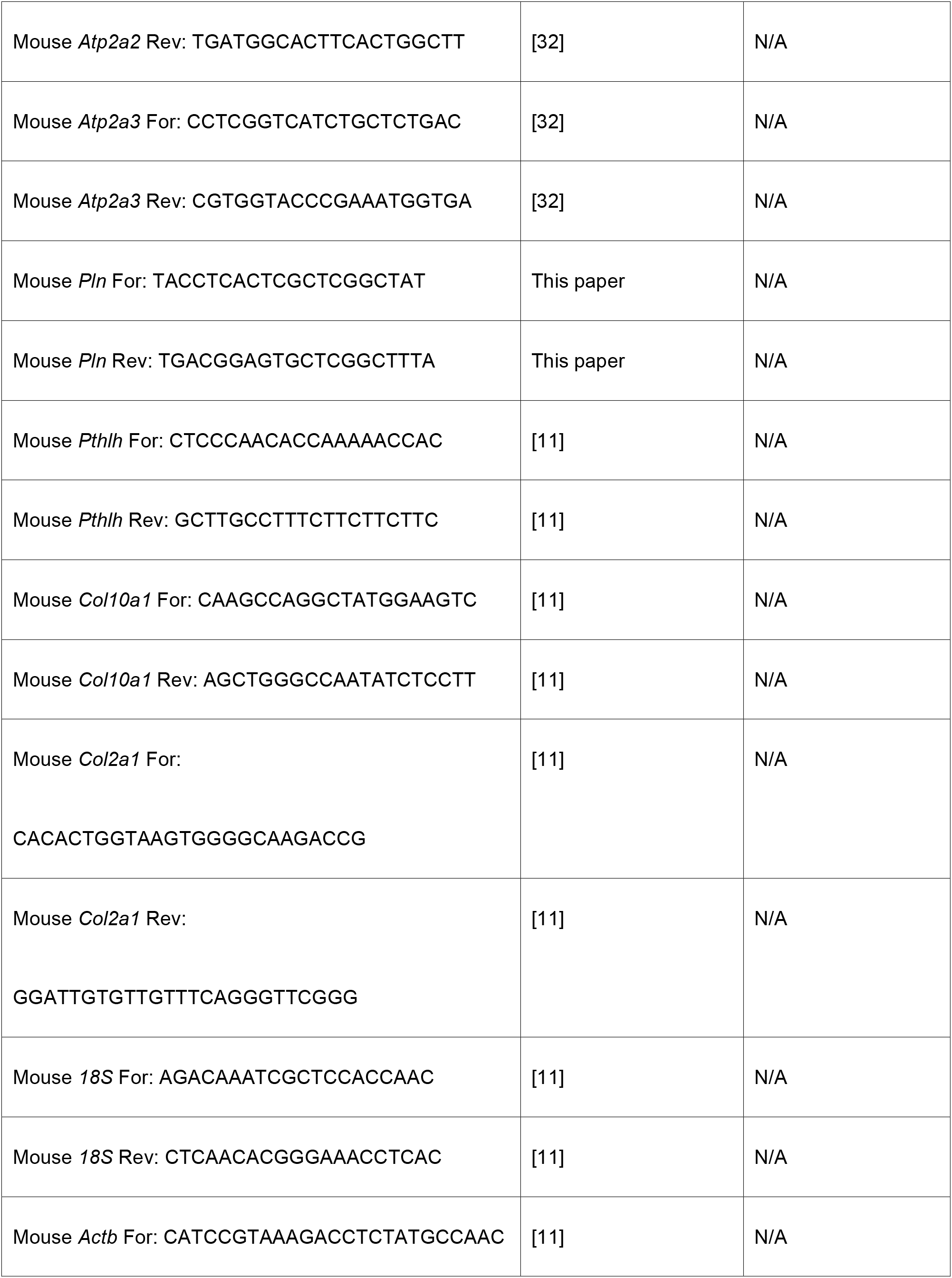
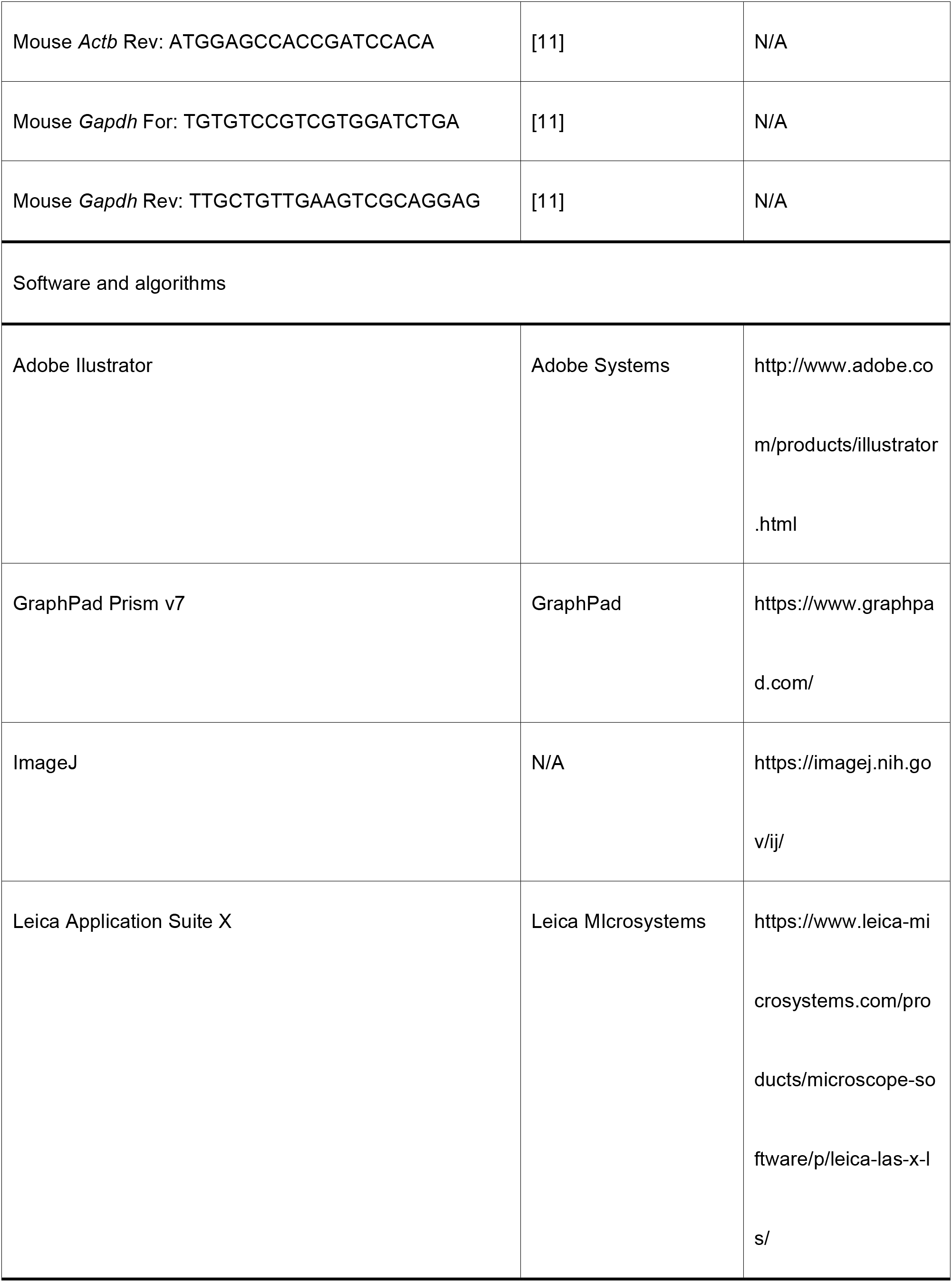

**Figure 6B-Source Data 1.**
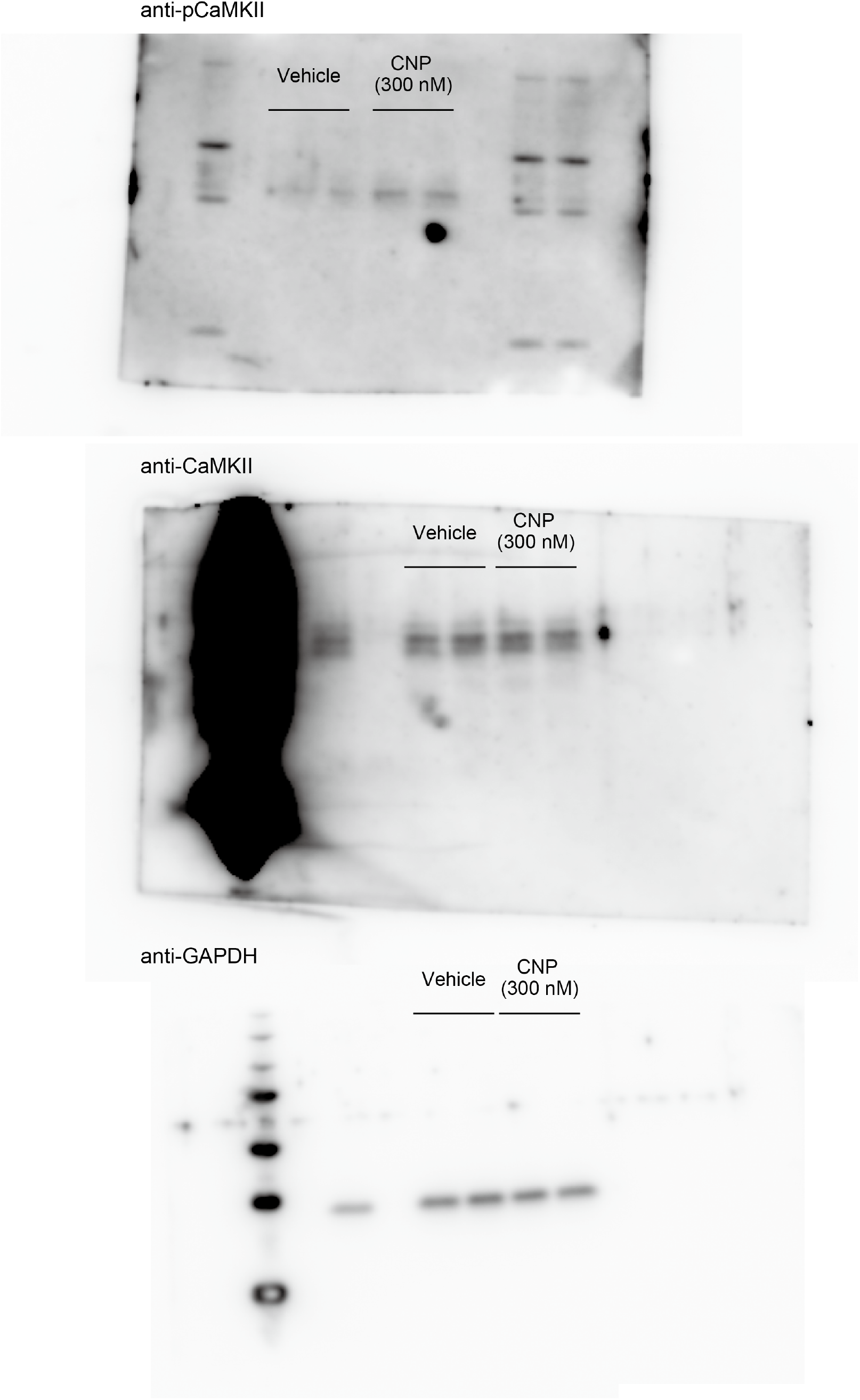
Data related to Figure 6B – Uncropped images.

**Figure 1-figure supplement 1B-Source Data 1.**
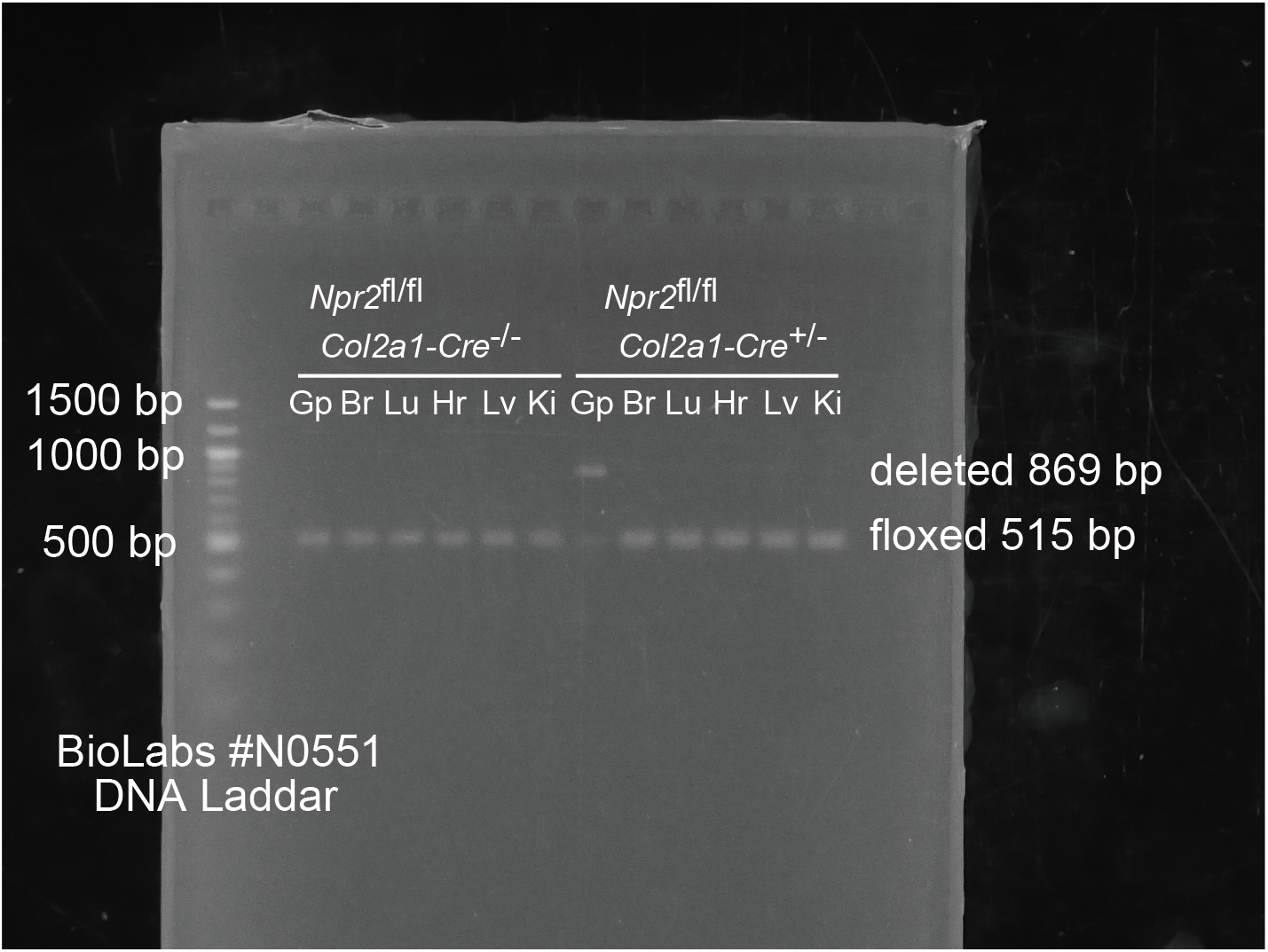
Data related to Figure 1-figure supplement 1B – Uncropped images.

**Figure 1-figure supplement 1B-Source Data 2.**
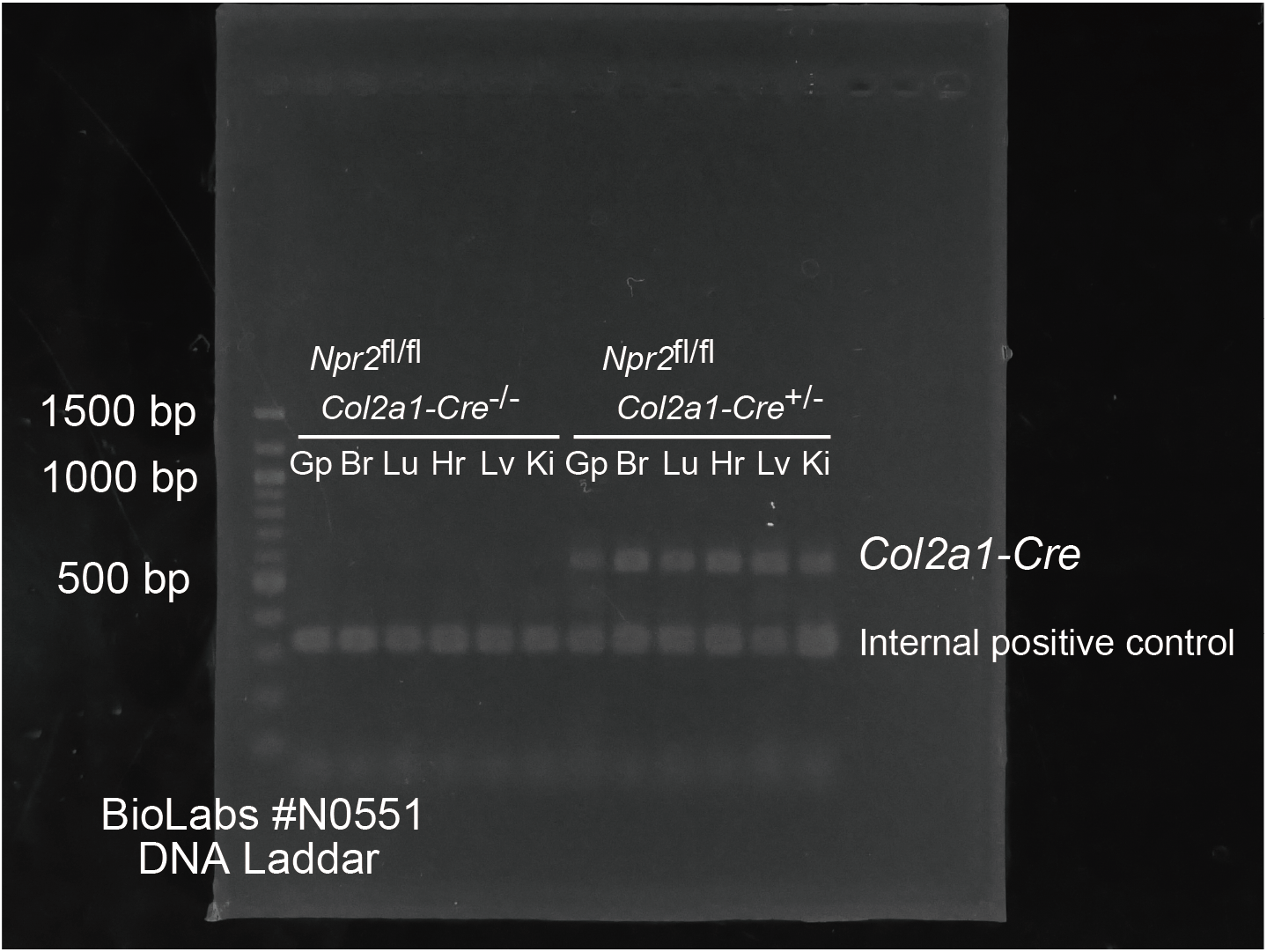
Data related to Figure 1-figure supplement 1B – Uncropped images.

